# Rapid intralacustrine evolution of an invasive pelagic three-spined stickleback (*Gasterosteus aculeatus*) ecotype in Lake Constance

**DOI:** 10.1101/2022.09.01.506194

**Authors:** Carolin Dahms, Samuel Roch, Kathryn R. Elmer, Albert Ros, Alexander Brinker, Arne Jacobs

## Abstract

The rapid invasion of the pelagic zone in Lake Constance by three-spined sticklebacks (*Gasterosteus aculeatus*) since 2012 and their subsequent drastic population growth has had stark ecosystem-wide effects, such as food-web shifts and declines in native biodiversity, including commercially important fish species. Yet, the origin of this invasive pelagic ecotype remains unclear. This study aims to determine if the pelagic ecotype arose *in situ* from the existing littoral population or following external colonisation, identify potential phenotypic differences between individuals from different habitats, and assess genomic signals of selection. Integrating RAD-sequencing of Lake Constance individuals and whole-genome sequence data for European outgroup populations, this study shows that the pelagic Lake Constance population likely arose recently within the lake from the littoral population, with only weak genome-wide differentiation between individuals from different habitats. This is further supported by minimal differences in meristic and morphometric traits, with shape differences only found between pelagic/inflow sticklebacks and littoral sticklebacks. Using genome scans, we identified multiple outlier loci between littoral and pelagic ecotypes across the genome, potentially suggesting early signs of sympatric speciation despite high connectivity. Furthermore, increased differentiation between pelagic and littoral sticklebacks for body shape-associated loci and the overlap of outlier loci with quantitative trait loci for ecologically relevant traits points toward a driving role of selection in this pelagic invasion. This study provides an important example of rapid ecological diversification from standing genetic variation and a rare case of littoral-pelagic ecotype divergence under high gene flow in a large oligotrophic lake. Ultimately, the results of this study will have major implications for the management of the invasive pelagic ecotype, and the entire stickleback population as a whole.

## Introduction

The introduction, establishment and spread of non-native species into novel habitats poses a serious threat to endemic biodiversity, ecosystem (Bax *et al*., 2003) and human health globally (Mainka and Howard, 2010). Freshwater systems have been particularly affected by the cascading abiotic and biotic effects of invasive species (Darwall *et al*., 2018), where, combined with other pressures such as climate change and pollution, the rate of species loss has exceeded those observed in terrestrial systems (Albert *et al*., 2021; Ricciardi and Rasmussen, 1999). While freshwater ecosystems harbour disproportionate amounts of biodiversity and provide essential ecosystem services (Darwall *et al*., 2018), these are threatened by anthropogenic activities, including intentional (e.g., for aquaculture or fishing enhancement) or accidental introductions of species presenting major pathways of freshwater invasions (Copp *et al*., 2005; Kiruba-Sankar *et al*., 2018).

Invasive species may harm native fauna and ecosystems indirectly by altering habitat conditions, such as habitat structure and the biogeochemical cycling (Crooks, 2002; Strayer, 2010); or directly through biotic interactions with cascading effects along the food chain (Gallardo *et al*., 2016). Biotic interactions include competition and predation that may hamper recruitment and decrease native species abundances (Bergström *et al*., 2015; Dudgeon, 2019; Roch *et al*., 2018). Other effects include the introduction of parasites and diseases (Cucherousset and Olden, 2011; Lymbery *et al*., 2014; Strayer, 2010), as well as reduced fitness and compromised genomic integrity through hybridization with related native species (Muhlfeld *et al*., 2009). Significant community-wide consequences might manifest also due to the evolutionary isolated and at times species-deprived states of freshwater systems (Cox and Lima, 2006), particularly in native peri-alpine and oligotrophic lakes where species may be vulnerable due to their lack of adaptation to invaders (Moyle and Light, 1996).

One such vulnerable system is Lake Constance, a large pre-alpine, oligotrophic lake in central Europe, where the inception of non-native three-spined stickleback (*Gasterosteus aculeatus*) has caused substantial ecosystem-wide changes (Gugele *et al*., 2020). While the exact origins of sticklebacks in Lake Constance are still under debate (Hudson et al., 2021), this small fish species was likely introduced around 150 years ago by deliberate releases into nearby streams and ponds by fishermen and aquarium hobbyists (Muckle, 1972). However, it was not until the middle of the 20th century that sticklebacks were first recorded in Lake Constance itself, where they subsequently spread throughout the nearshore of the lake within a few years (Roch et al. 2018) and since have been present in moderate abundance. This stickleback distribution drastically changed in late 2012 when a sudden massive increase in abundance and an expansion into the pelagic zone of Upper Lake Constance was documented (Alexander *et al*., 2016; Rösch et al. 2018; Eckmann & Engesser 2019); with an increase in small fish density from about 420 ± 145 (individuals ha^−1^, mean ± standard deviation) between 2009 and late 2011, to 2550 ± 800 between late 2012 and early 2014, and ultimately to 5300 ± 1970 from late 2014 to 2018 (Eckmann & Engesser 2019). Originally dominated by native whitefish (*Coregonus wartmanni*), the primary fisheries target in Lake Constance (Baer *et al*., 2017), the pelagic fish community experienced rapid change and constituted of sticklebacks by 95% after only two years (Alexander *et al*., 2016), with fisheries yields of whitefish declining by 50% in 2015 (Rösch *et al*., 2018). The loss of whitefish biomass and abundance is thought to be caused by sticklebacks being strong competitors for food such as zooplankton (*Daphnia* spp.; Bretzel *et al*., 2021, DeWeber *et al*., 2022); and by predation on whitefish eggs and larvae (Roch *et al*., 2018, Baer *et al*., 2021, Ros *et al*., 2019), which appear to lack adaptive predator avoidance compared to other lake species and consequently experience decreased growth and recruitment (Ros et al. 2019). Moreover, the increased pelagic stickleback abundance might cause further cascading effects through the food-web by shifting the species composition of pelagic zooplankton (Ogorelec *et al*., 2022), and changing densities and migration patterns of stickleback-feeding birds such as herons, kingfishers, and gulls (Werner *et al*., 2018). While these circumstances resemble those in the Baltic Sea, where a recent surge in native stickleback abundance has significantly affected the food chain and recruitment of native fish species (Bergström *et al*., 2015; Byström *et al*., 2015; Eklöf *et al*., 2020), such hyper-abundance of three-spined sticklebacks in a large oligotrophic lake is rare, even more so in pelagic waters, which represent an unusual habitat for this species (but see Erickson *et al*., 2016).

While Lake Constance three-spined sticklebacks provide an ideal opportunity to study the process and ecosystem-wide effects of freshwater invasions, this species has long been a model system for studying contemporary evolution across divergent ecological niches (Hudson *et al*., 2021). For example, the ability to rapidly adapt and genetically diverge in response to ecologically driven divergent selection in only few generations was demonstrated in lake and stream stickleback populations (Laurentino *et al*., 2020), which show substantial phenotypic variation in a range of traits such as foraging and defence performance (Arnegard *et al*., 2014; Lucek *et al*., 2014b; Schmid *et al*., 2019) and body size (Sharpe *et al*., 2008). Genetic evidence suggests that this rapid diversification across stream and lake niches was likely facilitated by introgression of multiple ancient Eastern and Western European lineages via secondary contact in the Lake Constance region, which facilitated the recruitment of adaptive alleles (Marques *et al*., 2016; Marques *et al*., 2019). In contrast to stream-like stickleback in other large, peri-alpine lakes, such as Lake Geneva, Lake Constance stickleback, which likely first established in littoral waters in the 1950’s, have a large-bodied, fully plated pelagic phenotype (Hudson *et al*., 2021). Lake Constance stickleback largely show Baltic Sea ancestry, and it has been hypothesised that this Eastern European ancestry, resulting in pelagic phenotypes, might have promoted the recent population expansion from the littoral to the pelagic zone (Hudson *et al*., 2021; Lucek *et al*., 2014a). However, so far there is no clear evidence for drivers that could explain such an expansion (Baer et al. 2022, Ogorelec et al. 2022). Similarly, stream-adaptive alleles from Western European stickleback lineages are thought to have promoted the colonisation of stream habitats in Lake Constance (Marques *et al*., 2019). In the case of the pelagic niche expansion in Lake Constance, it remains untested whether the open-water stickleback populations i) have evolved in sympatry from the littoral counterparts, ii) if littoral and pelagic populations form genetically distinct groups, and iii) if this ecological divergence has been facilitated by recent colonisation and hybridization. In either scenario, this system could provide a unique opportunity to study ecological speciation of a freshwater invader in its early stages.

Given their dramatic effect on the lake’s ecosystem and fishery, this study aims to gain better understanding of the genetic and phenotypic differences and evolutionary origins of the recently emerged pelagic three-spined stickleback population in Lake Constance. Using Restriction-site Associated DNA (RAD) sequences and whole genome data from outgroup populations, we first investigated whether the pelagic population is genetically distinct from the littoral and tributary (here referred to as ‘inflow’) sticklebacks. Populations from tributaries were included in the current study to test the hypothesis that pelagic sticklebacks migrate to tributaries for spawning, promoting reproductive isolation from the littoral ecotype. Population genetic analyses were complemented by investigations on morphometric and meristic traits. Second, we aimed to identify genetic regions underlying adaptations to the pelagic habitat by searching for genetic differences potentially under selection between littoral and pelagic sticklebacks, and identifying genomic regions associated with ecologically relevant traits in this study and previous quantitative trait mapping studies (Erickson *et al*., 2016; Peichel and Marques, 2017). Although the pelagic expansion is only a decade old (5-7 stickleback generations), rapid evolution in novel environments in just a few generations has been demonstrated in stickleback (Lescak *et al*., 2015). In this study system, genetic differentiation could be a possible result of rapid alternative sorting of adaptive alleles into littoral and pelagic ecotypes. The divergent lifestyles (such as foraging and defence strategies) between benthic and limnetic habitats could produce strong divergent selection pressures. Overall, this study provides important insights into the potential of an invasive species to rapidly colonise novel habitats and the underlying genetic architecture, which will directly inform the management of stickleback in Lake Constance.

## Material and Methods

### Sampling

Sampling took place in Lake Constance, which at a size of about 535 km^2^ and an average depth of 100 m, is one of the largest lakes in Europe (Petri, 2006). Sampling in the Upper Lake of Lake Constance was carried out in spring and summer 2019 and three different methods were utilised: fish were caught by trawling in the pelagic zone, gillnetting in the littoral zone, and electrofishing in three tributaries of Lake Constance (Fig. 1; Table 1). Details on the respective fishing method can be found in Gugele *et al*. (2020). A total of 95 sticklebacks were sampled. Fish from trawling and electrofishing were euthanized with clove oil (1 ml/L water, Euro OTC Pharma, Bönen, Germany) and total length (accuracy: 0.1 cm) was recorded. Each individual was then photographed laterally using a digital camera (Pentax K3 II, with 18-135 mm lens and fixed focal length). For the genetic analysis, a piece of the caudal fin (5pprox.. 0.5 cm²) was fixed in pure ethanol. The sex was determined by dissection and the fish were stored at −20 °C until further processing. Fish sampling was carried out according to local regulations (“Landesfischereiverordnung Baden-Württemberg”, LfischVO).

### Meristic and morphometric traits

To determine the number of lateral bone plates, sticklebacks were stained according to a protocol modified from Peichel *et al*. (2001). Individuals were placed in neutral buffered 10% formalin (Sigma-Aldrich, St. Louis, USA) for one week and subsequently rinsed in ultrapure water (0.2 μm pore size, Arium 611, Sartorius Stedim Biotech, Göttingen, Germany) for 24 h. For staining of the bony structures, individuals were placed in 0.008% alizarin red (alizarin red S monohydrate, Waldeck Chemie, Münster, Germany) and 1% potassium hydroxide (EMSURE, Merck, Darmstadt, Germany) for 24 h. For subsequent decolourisation, individuals were placed in 1% potassium hydroxide and 1% hydrogen peroxide (CHEMSOLUTE, Th.Geyer, Renningen, Germany) solution for five days. Finally, sticklebacks were fixated in 48% ethanol for 24 h and then stored in 70% ethanol. The number of plates was determined for both sides of each individual under a dissecting microscope (Zeiss, Stemi SV6, 8x – 50x magnification) and the mean number of both sides was used for further analysis (Fig. 2A). All sticklebacks were classified according to Bell (2001) into “fully plated” (>30 lateral plates), “partially plated” (<34 and >10 lateral plates, with a gap) and “low plated” (<10 lateral plates). The statistical comparison of lateral plate number between the individual sampling sites was done using a pairwise Steel-Dwass test (Steel 1960). Finally, differences in total length of sticklebacks from each sampling site was tested using an ANOVA and a Tukey-Kramer HSD *post hoc* test (Sokal and Rohlf 1995). Both tests were conducted in JMP 16.0.0 (64 bit, SAS Institute, Cary, USA).

**Table 1:** Summary of individual sampling events and sampling success in Lake Constance. n = number of individuals sampled

| Sampling site | Sampling date | Sampling method | Habitat | n<br>total | n<br>female | n<br>male |
| --- | --- | --- | --- | --- | --- | --- |
| Pelagic zone | 26.03.2019 | Trawling | Pelagic | 32 | 13 | 20 |
| Littoral zone | 04.06.2019/<br>05.07.2019 | Gillnets | Littoral | 32 | 19 | 13 |
| Alter Rhein | 16.05.2019 | Electrofishing | Inflow | 11 | 7 | 4 |
| Nonnenbach | 08.04.2019/<br>17.04.2019 | Electrofishing | Inflow | 10 | 6 | 4 |
| Brunnisach | 10.04.2019/<br>17.04.2019 | Electrofishing | Inflow | 10 | 6 | 4 |

**Fig. 1.**
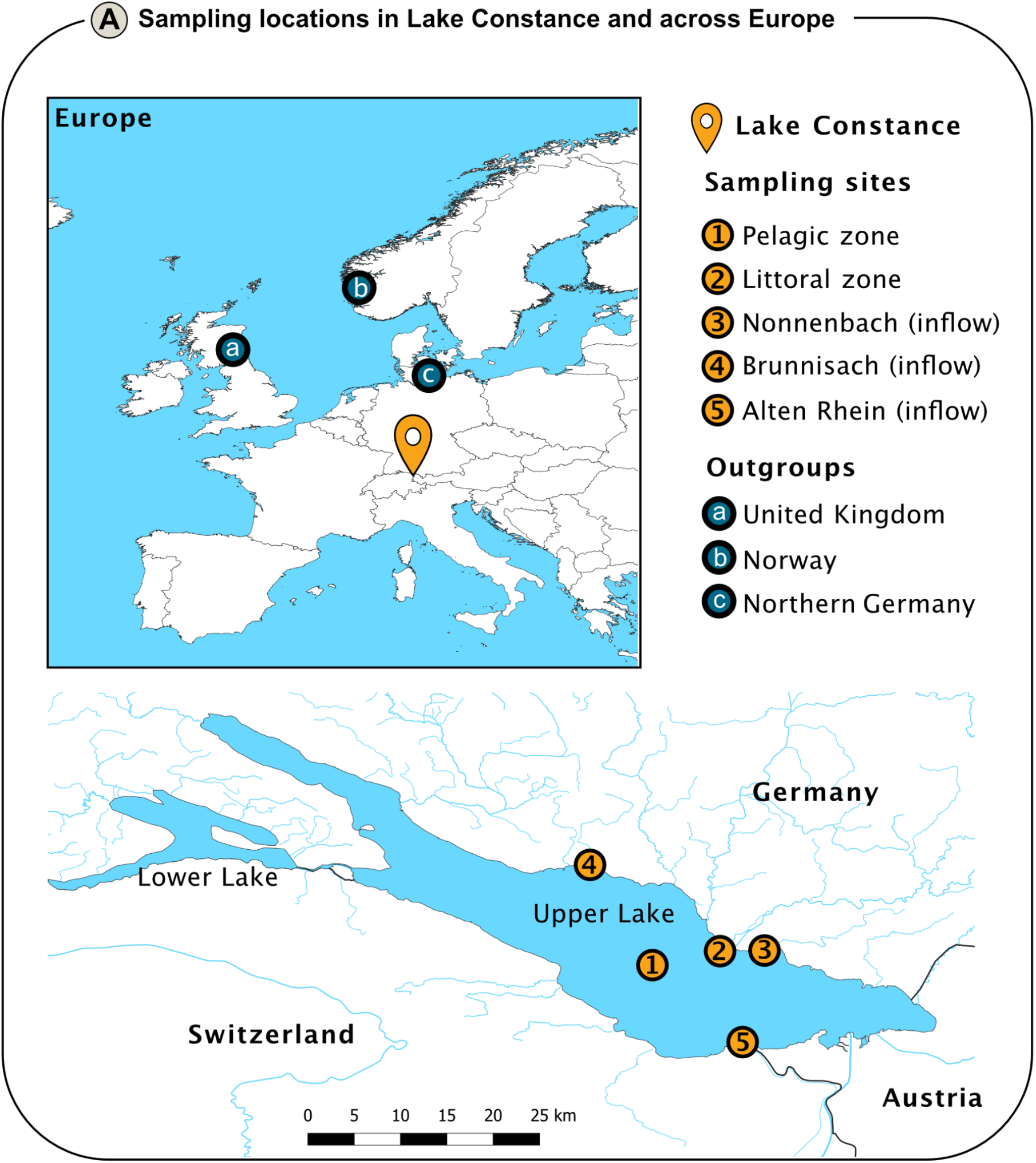
Overview map showing the position of the individual sampling locations in Lake Constance and locations of outgroup populations (insert) used in the genetic analysis. Population information and geographic coordinates for all samples are provided in Supplementary Data S1.

**Fig. 2.**
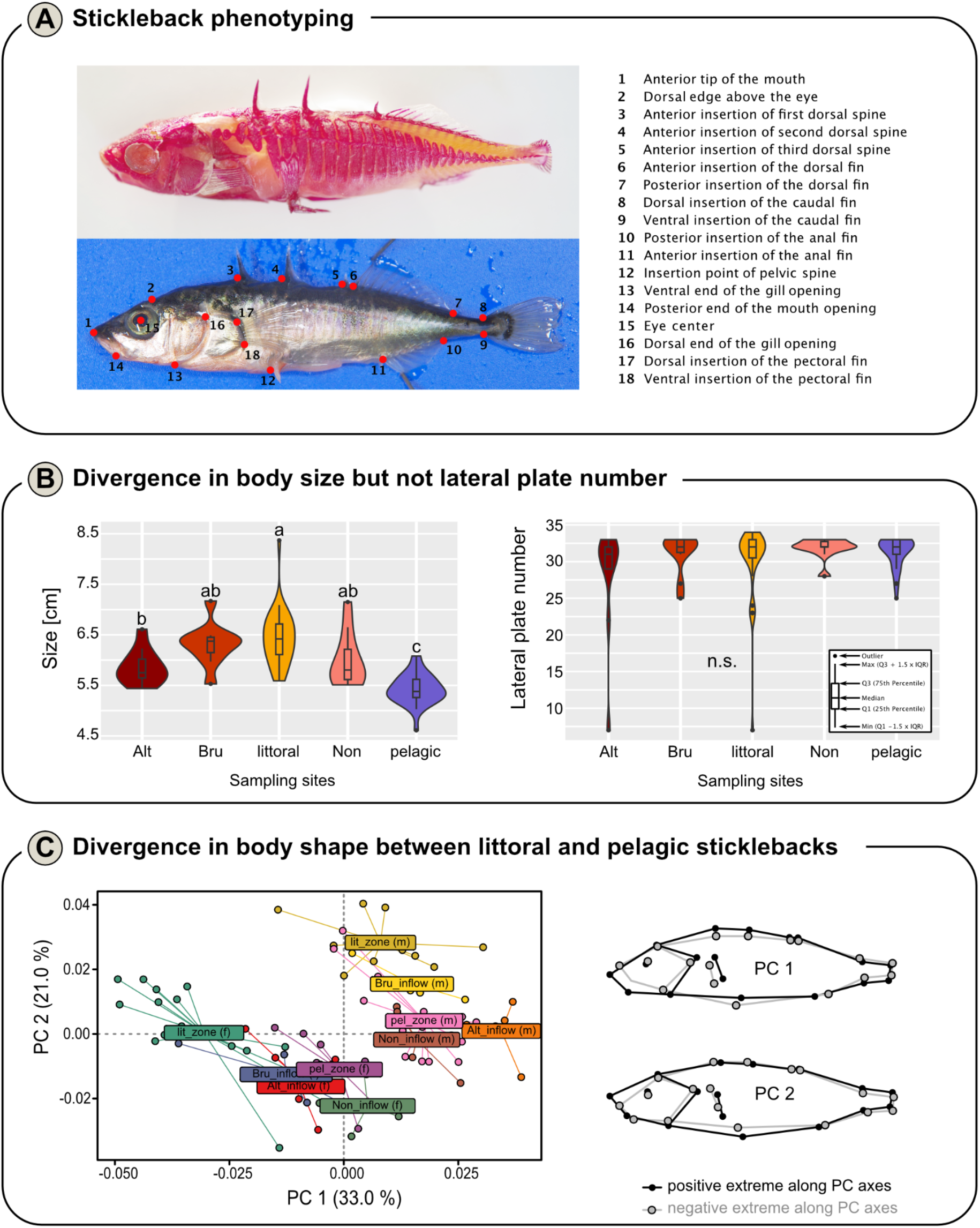
Phenotypic diversity of Lake Constance sticklebacks. **(A)** (Top) Stickleback with stained bony structures (for more details, please refer to the text), allowing the determination of the lateral plate number. (Bottom) Position of 18 unique reference points (“landmarks”) on the body for shape analysis. (Right) Description of the individual locations of the landmarks. **(B)** Combined boxplot and violin plot to illustrate variation in size (left) and lateral plate number (right) of sticklebacks from different sampling sites. Lower case letters indicate statistically significant differences (size: ANOVA + Tukey-Kramer HSD post hoc test, lateral plate number: pairwise Steel-Dwass test). Box plots defined in the insert panel on the right **(C)** Scatterplot showing the first two principal components (PCs), which explain most of the variance of the data (see axis labelling in percent). Sticklebacks were grouped according to sampling site (Bru = Brunnisach, Non = Nonnenbach inflow, Alt = Alter Rhein inflow) and sex (m = male, f = female). Wireframe graphs of the shape changes along the first two PCs in the PCA are shown on the right.

For the morphometric analysis, 18 unique reference points („landmarks”) were placed on digitale images of sticklebacks using TPSDig 2 (V2.31, James Rohlf, Stony Brook University, New York, USA) (Fig. 2A). Statistical analysis was performed using the R package “geomorph” (V4.0.3; Baken *et al*., 2021). A Generalised Procrustes Analysis (GPA; Gower, 1975) was performed using the “gpagen” function to remove differences in size, position, and orientation. Possible errors in landmark placement were identified using the “plotOutliers” function and affected individuals were excluded from further analysis as appropriate. In a next step, a model was developed which took into account fish size (i.e. allometric effects) as a covariate, sampling site as a fixed factor and sex as a random factor, nested within the fixed factor: *coords = log(size) + sex + sex:site*. A permutation-based Procrustes ANOVA using residual randomization (function: “procD.lm” and “anova”, permutations = 9999; estimation method: ordinary least squares; Goodall, 1991) was used to examine which factors have a statistically significant effect on shape. A second permutation-based Procrustes ANOVA (permutations = 9999; estimation method: ordinary least squares) was used to test if allometric effects were common for all individuals or unique for each sampling site or sex. Pairwise *post hoc* test of sampling sites was performed with the “pairwise” function of the R package “RRPP” (V1.2.3; Collyer and Adams, 2018; Collyer and Adams, 2021) and based on the previously developed model and a null model considering only size and sex (permutations = 9999, test type = distance between vectors (“dist”), confidence = 0.95). The obtained p-values were adjusted using the Bonferroni-Holm method (Holm, 1979). The Procrustes shape coordinates of each individual were used in a principal coordinate analysis (PCA; Sokal and Rohlf, 1995) to illustrate shape differences between sex and sampling sites respectively. Principal components (PCs) were calculated using the “gm.prcomp” function of the “geomorph” package and the first two PCs, explaining the greatest variation, were plotted. Additionally, shape changes along the two PCs were illustrated in a wireframe graph. All calculations were performed, and figures were generated using R Version 4.1.3 (R Core Team, 2022) and RStudio Version 2022.02.3 (RStudio Team 2020).

### DNA extraction and RAD sequencing

DNA for RAD sequencing was ultimately extracted from frozen brain tissue as samples from fins of these fish yielded low quality DNA. DNA extraction was carried out according to the manufacturer’s instructions. Approximately 15 mg frozen brain tissue was added to a 1.5 mL tube containing a buffer and two 2.4 mm diameter metal beads (OMNI international, Kennesaw, USA). Tissue was homogenised for three seconds at maximum speed (Bead Ruptor 4, OMNI international, Kennesaw, USA). The homogenate was digested for three hours at 55 °C with continuous mixing (150 RPM) on a thermomixer. RNA was degraded by incubation with RNAse for one hour, and thereafter DNA was extracted from the homogenate using the provided buffers and spin columns provided in the kit. DNA was eluted in the spin column with 50 μL of elution buffer yielding 35 – 160 ng/μL DNA (NanoDrop 2000c, Thermo Fisher Scientific, Waltham, USA). Only samples that showed none or weak degradation, as determined using agarose gel electrophoresis, were selected for NGS. Sample concentrations were standardised to 20 ng/μL, and final concentration of DNA for NGS was checked using fluorometry (Qubit 3, Life Technologies, USA). These samples were shipped to Floragenexs (Eugene, Oregon, USA) for RAD library preparation and sequencing. Restriction-site Associated DNA Sequencing (RAD-seq) libraries were generated as detailed in Etter *et al*. (2011) using the restriction enzyme *SbfI* (New England Biolabs).

### Outgroup populations

Whole genome sequences of 30 individuals representing outgroups from the Baltic and North Sea were acquired from published studies (Feulner *et al*., 2015; Jones *et al*., 2012b). Four whole genome sequences of a sister species *Gasterosteus nipponicus* (Yoshida *et al*., 2014), which is estimated to have diverged from *G. aculeatus* approximately 0.68 million years ago (Kitano *et al*., 2009), were retrieved as an outgroup for phylogenetic analyses. Population information, geographic coordinates and ecotypes of all samples are provided in Supplementary Data S1.

### Sequencing and SNP calling

FASTQ data files provided by Floragenex were demultiplexed using GBSX v.1.3 (Herten *et al*., 2015). Reads were mapped to the *G. aculeatus* reference genome (release 92), retrieved from the Ensembl database (Hunt *et al*., 2018), using BWA v.0.7.17 (Li and Durbin, 2010). Any reads that included bases with a Phred quality score < 20 were removed. Genotypes were hard-called (i.e., determined according to the highest probability) using Stacks v.2.6 *gstacks* module (Catchen *et al*., 2013). The Stacks output was subsequently converted into a VCF (Variant Call File) which was filtered using VCFtools v.0.1.16 (Danecek *et al*., 2011). The filtering removed indels (*--remove-indels*), SNPs with minor allele frequencies below 0.05 (*--maf 0*.*05*) and more than 25% missing data (*--max-missing 0*.*75*). Only loci with a minimum read depth of 5 (*-- minDP 5*), a minimum mean depth of 5 (*--min-meanDP 5*), a maximum depth of 50 (*--maxDP 50*), a maximum mean depth of 50 (*--min-meanDP 50*) were kept, and variants with quality below 10 (*--minGQ 10*) were removed. This procedure produced a VCF with 28,194 out of originally 361,485 variant sites.

### Population genetic structure

Geographic structure across all Lake Constance sites was explored by performing a Principal Component Analysis (PCA) on the filtered SNP data set using ade4 v.1.7-16 (Dray and Dufour, 2007). For the analysis, SNP data were pruned for linkage disequilibrium in Plink v.1.9 (Purcell *et al*., 2007) in 25 SNP windows with a five SNP window shift, and an r^2^ of 0.5, reducing the original dataset from 28,194 to 11,184 sites. The sex chromosome 19 was removed to avoid any sex-based bias in down-stream analyses (Peichel *et al*., 2004).

Individual ancestry was assessed using Admixture v.1.3.0 (Alexander *et al*., 2009), using the LD-pruned SNP dataset. Clusters (K) = 1-7 were tested to determine the optimal number of K by calculating the tenfold cross-validation standard errors in Admixture.

### Genetic differentiation

Genome-wide distribution of genetic differentiation was studied by calculating haplotype-based relative allelic differentiation (*F*_ST_*’*) and absolute divergence (*D*_*xy*_) between population pairs using the *populations* module (*--fstats*) in Stacks for all loci containing filtered SNPs. Haplotype-based estimates have the advantage of accommodating loci with more than two alleles contrary to SNP-based statistics (Nei, 1973). Unlike *F*_ST_*’, D*_*xy*_ does not depend on contemporary levels of interpopulation diversity (Charlesworth, 1998; Charlesworth *et al*., 1997; Jakobsson *et al*., 2013; Nei, 1973), but is affected by levels of diversity within the ancestral population (Cruickshank and Hahn, 2014). Gene diversity (*Hs*), a haplotype-based equivalent to nucleotide diversity which is corrected for sampling bias originating from sampling small sample sizes (Nei, 1987), was also calculated with *populations* across the loci containing the 28,194 SNPs previously filtered with VCFtools.

To identify genomic regions potentially under selection in pelagic Lake Constance sticklebacks, z-transformed *F*_ST_’ and *D*_*xy*_ estimates for the pelagic-littoral population comparison were computed. Loci with z-transformed *F*_ST_’ ≥ 3 were classified as differentiated outlier loci, and those also with z*D*_*xy*_ ≥ 3 were classified as putatively under selection. A z-transformed value ≥ 3 approximately corresponds to a p-value below 0.01.

Identified outlier loci were further tested for potential selection by comparing their interpopulation gene diversity differences (Δ*Hs = Hs*_*pelagic*_ *- Hs*_*littoral*_) to the genome-wide background. The expectation was that Δ*Hs* would be higher in loci under selection in the pelagic populations compared to the genomic background, driven by reduced *Hs* in the pelagic ecotype. The data was tested for any deviations from normality and was found to violate the assumption of homoscedasticity, as suggested by a Levene’s test finding significantly uneven variances between Δ*Hs*_*outlier*_ and Δ*Hs*_non-outlier_ (*F*_*1*,16436_ = 881.52, *p* < 0.001). Therefore, medians were compared with a non-parametric two-sided Wilcoxon rank sum test. While the analyses involved primarily the littoral-pelagic comparison which was the main focus of this study, genetic differentiation estimates among the remaining populations and outlier loci overlaps among all pair-wise population comparisons are detailed in Supplementary Fig. S2 and Supplementary Fig. S3, respectively.

### Genome-wide association analysis

Genetic association mapping using Genome-wide Efficient Mixed Model Association (GEMMA v.2.1; Zhou et al., 2013) was conducted to identify genome-wide signals of association between markers and two phenotypes: body shape and plate number in the 95 Lake Constance sticklebacks. Associations with total length were not performed, as fish were sampled at different times of the year and might therefore differ in age and size.

We fitted standard Bayesian Sparse Linear Mixed Models (BSLMM) for each phenotype, providing the same genotype and relatedness matrix input files. The BSLMM is well suited when the genetic architecture of a trait is unknown, as it combines the benefits from linear mixed models with sparse regression models (Armstrong et al. 2018). However, as the BSLMM does not allow covariate files to correct for sex, we corrected traits that showed sex-differences (body shape) using a linear model in R. We used the residual body shape PC scores from these linear models as input for the BSLMM analyses. We fitted five separate BSLMMs for each phenotype to account for differences between runs. For each of the five runs per phenotype, the results were first averaged across all chains and subsequently across runs. In contrast to standard linear mixed models, the BSLMM also estimates a range of hyperparameters describing the genomic architecture of a trait, such as the proportion of variance in phenotypes explained by all SNPs (PVE), the proportion of variance explained by sparse effect loci (PGE), and the number of variants with major effects (n gamma). We estimated the means, median, and 95th confidence interval (CI) for all these parameters, first within replicated runs and then averaged them across runs. Furthermore, we identified SNPs as those with an average posterior inclusion probability (PIP) above 0.01 as significant (Comeault et al. 2014), and those with PIP > 0.1 as highly significant (Chaves et al. 2016).

Furthermore, we tested if SNPs that show significant associations with phenotypes were also significant outlier loci or showed increased genetic differentiation between littoral and pelagic sticklebacks, which would suggest potential selection acting on these phenotypes. To test if phenotype-associated SNPs showed increased genetic differentiation and divergence compared to a random genomic background, we performed random resampling of the same number of SNPs from the entire SNP datasets, estimated the mean *F*_*ST*_*’* and *D*_*xy*_ for the corresponding haplotype, and repeated this 10,000 times to create a null distribution. Subsequently, we compared the means *F*_*ST*_*’* and *D*_*xy*_ of the phenotype-associated SNPs and the null distribution using a Wilcoxon test. We did this for each phenotype, and for the sex-chromosome and autosomes separately.

### Association with quantitative trait loci

To investigate whether the identified outlier loci were potentially associated with phenotypic traits in *G. aculeatus*, we searched for overlaps between outlier loci (*zFst’* ≥ 3 and *zD*_*xy*_ ≥ 3) and an extensive list of genomic regions underlying adaptive phenotypes previously identified by Quantitative Trait Locus (QTL) mapping studies (Peichel and Marques, 2017; see Supplementary Data S2). These 28 studies addressed divergence in a range of morphological, physiological and behavioural traits between marine and different freshwater ecotypes (benthic, limnetic, lake and stream) in three-spined sticklebacks (Albert et al., 2008; Arnegard *et al*., 2014; Berner *et al*., 2014; Cleves *et al*., 2014; Colosimo *et al*., 2004; Conte *et al*., 2015; Coyle *et al*., 2007; Cresko *et al*., 2004; Ellis *et al*., 2015; Erickson *et al*., 2016; Glazer *et al*., 2014; Glazer *et al*., 2015; Greenwood *et al*., 2015; Greenwood *et al*., 2011; Greenwood *et al*., 2013; Kimmel *et al*., 2005; Kitano *et al*., 2009; Liu *et al*., 2014; Malek *et al*., 2012; Miller *et al*., 2007; Miller *et al*., 2014; Peichel *et al*., 2001; Rogers *et al*., 2012; Shapiro *et al*., 2004; Wark *et al*., 2012; Yong *et al*., 2016; Yoshida *et al*., 2014). QTL coordinates were extracted according to the BROAD S1 genome assembly, with corrections for testing QTL region overlaps (Peichel and Marques, 2017).

Only a third (101) of the associated QTLs originated from studies focusing on benthic-limnetic divergence, and majority of QTLs (178) originate from different freshwater and marine systems (Supplementary Data S2). The rationale for retaining QTLs from systems different to Lake Constance in the analyses was that although causative mutations across populations might differ, the same genes could be associated with the divergent traits observed in Lake Constance sticklebacks. QTL regions with unmapped chromosomes were excluded.

Overlaps between RAD-seq loci and QTL were estimated following Marques et al. 2016. In brief, QTL regions were defined either based on 95% confidence intervals, if reported, or based on the peak marker ± 1Mb. Furthermore, we added ± 10kb to SNP positions as a buffer to account for the low density of SNPs from RAD sequencing.

### Phylogenetic analyses

To further infer phylogenetic relationships including stickleback populations from Northern Europe, a maximum likelihood tree with migration events among *Gasterosteus* populations from Lake Constance and Great Britain, Norway and Northeast Germany was reconstructed. For this analysis TreeMix v.1.12 was used, which utilises allele frequencies and a Gaussian approximation accelerating inference of genetic drift (Pickrell and Pritchard, 2012). To maximise the amount of data from a combined RAD-seq and whole genome data set, the input file for this analysis was obtained from genotype likelihoods calculated in ANGSD v.0.934 (Korneliussen *et al*., 2014). First, a list of sites with quality control (*-uniqueOnly 1, -remove_bads 1, -minQ 20, -minMapQ 20, -SNP_pval 1e-6*), a minor allele frequency of 0.05 (*-minMaf* 0.05) and allowing for 50% missingness (-minInd 64) was used to calculate minor allele frequencies for each population, which were filtered further for 20% missingness and combined into a final data set of 11,896 sites.

The TreeMix analysis was then run as per Dahms et al. (2022). Briefly, 500 TreeMix bootstrap replicates were created using a SNP block size of 100 for estimating the covariance matrix without migration events. *G. nipponicus* was indicated as the outgroup (*-root*) in the trees, from which a consensus tree was obtained in PHYLIP Consense v.3.697 (Felsenstein, 2005). To determine the most likely number of migration events (*m*), TreeMix was run with *k* at 100, the prepared consensus tree provided as the starting tree (*-tf*), from which a range of migration events (from one to 10) with 10 iterations for each *m* was tested. Model likelihoods for each number of migrations were compared with the R package OptM v.0.1.3 (Fitak, 2018), which suggested an optimum *m* of four based on model likelihoods. Lastly, 30 independent runs, each starting from a random seed and earlier obtained consensus tree, were performed with 4 migration events and *G. nipponicus* set as the root. The final tree was chosen according to highest likelihood and plotted with node bootstrap values indicating relative confidence in the tree topology. Bootstrap values were calculated as the percentage of bootstrap replicates (n = 500) containing the node position chosen in the consensus tree. The least significant p-value from all runs, and migration weights (i.e., the percentage of ancestry received) averaged over 30 final runs for each migration event are provided in Supplementary Table S4. Plotted residual and drift matrices can be found in Supplementary Fig. S4.

All figures produced in this study were plotted in R v.4.0.2 (R Core Team, 2013) and R Studio v.1.3.1073 (RStudio Team, 2019).

## Results

### Meristic and morphometric traits

There were statistically significant differences apparent when comparing total length of sticklebacks between sampling sites (ANOVA, F_4,90_ = 23.1534, P < 0.0001; Fig. 2B), with pelagic individuals being smaller than those in all other groups. When looking at the lateral plate number, no differences between sampling sites were found (Steel-Dwass test, P > 0.05, Fig. 2B). Of all 95 sticklebacks examined, 77.9% were fully plated, 20.0% partially plated and 2.1% low plated.

The morphometric analysis revealed that size had a statistically significant effect on the shape of examined sticklebacks, indicating allometric effects (ANOVA, P < 0.0001; Table S1. These allometric effects were common to all sampling sites and both sexes studied (Table S2). Furthermore, sex and the sampling site had a statistically significant influence on the shape of the fishes after correcting for allometry (ANOVA, sex: P < 0.0001, sampling site: P < 0.0001; Table S2). Pairwise comparison of the sampling sites showed that the shape of fish from the littoral zone was statistically significantly different from all other sites (Table 1). The results of the performed PCA are shown in Fig. 2C, where the first two principal components accounted for 54% of the variance in the data. Changes in shape (Fig. 2C) were mainly evident in the head region, the positioning of the pectoral fin, as well as in the general body contour.

*statistically significant after Holm-Bonferroni correction (α = 0.05; Holm 1979)

**Table 1.** Results of the pairwise comparison of the shape of sticklebacks from different sampling sites (littoral = littoral zone, pelagic = pelagic zone, Inflow = Nonnenbach, Brunnisach, Alten Rhein). **Upper triangle:** pairwise procrustes distances between means. **Lower triangle:** pairwise p-values between means.

|  | Alten Rhein | Brunnisach | Littoral | Nonnenbach | Pelagic |
| --- | --- | --- | --- | --- | --- |
| Alten Rhein |  | 0.01739166 | 0.03533241 | 0.01404069 | 0.01253745 |
| Brunnisach | 0.707 |  | 0.02146350 | 0.02201050 | 0.01951720 |
| Littoral | 0.001* | 0.020* |  | 0.04093899 | 0.03781088 |
| Nonnenbach | 1.000 | 0.146 | 0.001* |  | 0.01256773 |
| Pelagic | 1.000 | 0.707 | 0.001* | 1.000 |  |

\*statistically significant after Holm-Bonferroni correction ( $\alpha = 0.05$ ; Holm 1979)

### Weak population genetic structuring

The admixture analysis suggested that sticklebacks in Lake Constance are highly admixed, as the genetic structure was best explained with 1 cluster (K = 1; Fig. S1). High admixture and low population structure was also supported by the PCA analysis (Fig. 4). Population PC centroids suggested some degree of clustering, with littoral and pelagic populations grouping closer compared to inflow populations, however PC1 and PC2 only explained negligible variation in genetic structure among individuals (1.75% and 1.52%, respectively).

Furthermore, genetic differentiation was low between all Lake Constance populations, both in relative divergence (mean *F*_ST_*’* = 0.005 ± 0.002) and absolute divergence (mean *D*_*xy*_*’* = 0.001 ± 2.61 × 10^−5^) (Table S3). Differentiation between the pelagic and littoral three-spined sticklebacks was the lowest among all pairwise comparisons (*F*_ST_*’* _LIT-PEL_ = 0.002).

### Evolutionary origin of pelagic stickleback in Lake Constance

To infer the evolutionary origin of pelagic Lake Constance stickleback, we reconstructed phylogeny of stickleback from Lake Constance and additional European locations (Fig.1). The phylogenetic relationship between Lake Constance stickleback and outgroup populations was best explained by a maximum likelihood tree in TreeMix with four migration events [ln(likelihood) = 421.377] (Fig. 3B). The final tree explained 99.5% of the variance in relatedness between populations and placed all Lake Constance populations into a monophyletic clade. Confidence in tree topology was generally high (90-100%) with few exceptions; no bootstrap values were reported for some nodes, such as in the Lake Constance clade, where TreeMix slightly changed the topology of the consensus tree (Fig. 3B). The tree placed the oldest lineages in the North Sea, followed by the Baltic Sea, among which freshwater populations were suggested to have experienced admixture – e.g., between British Tyne River (TY_F) and a northern German river population (GER_R), and Norwegian river (NOR_R) and lake populations (NOR_L). The longest tree branch representing high levels of drift branched the Lake Constance populations off from the Baltic lineage (Fig. 3B). While very low levels of drift were observed among Lake Constance populations, the pelagic (PEL) population was suggested to be the most divergent in this clade (Fig. S2). The four most likely migration edges inferred by TreeMix were all statistically significant (p < 0.05). Admixture events had generally low ancestry percentages received from source populations (i.e., migration weight *w*), ranging from 0.03 to 0.125, except between NOR_R and NOR_L (*w* = 0.32 ± 0.03; Table S4).

**Fig. 3.**
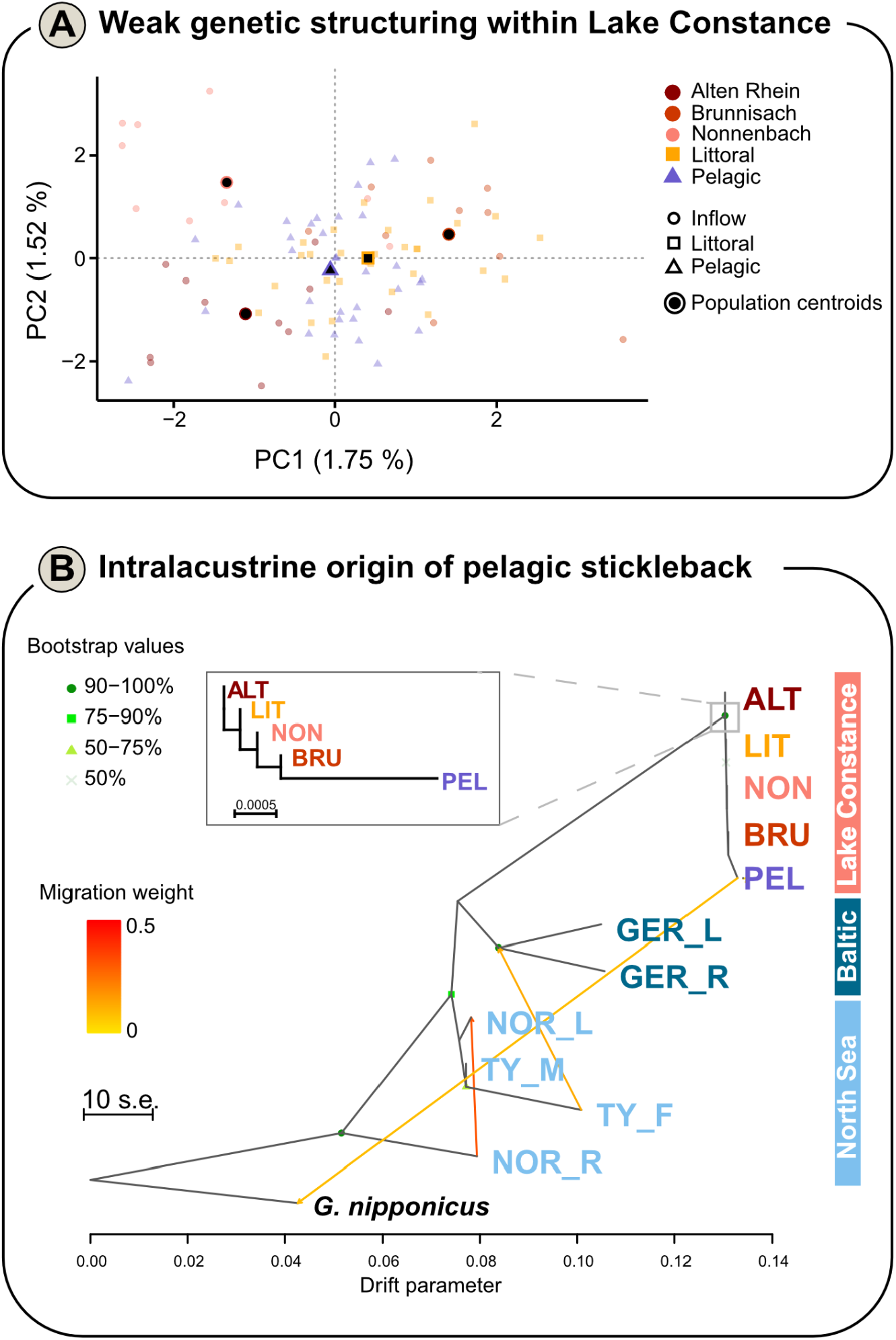
Population structure and evolutionary origin. (**A**) Principal Component Analysis representing individual structuring across Lake Constance ecotypes and populations. Analysis was performed on pruned data excluding the sex chromosome (11,184 SNPs). Colours indicate different sampling sites, while shapes represent ecotypes – inflow (circle), pelagic (square), pelagic (triangle). Smaller, lighter data points show individual variation, while the bigger shapes with a black centre indicate population Principal Component centroids, which were calculated as the mean of both the 1^st^ and 2^nd^ axes. (**B**) Maximum likelihood tree from TreeMix for Lake Constance and outgroup populations. Population names are coloured according to location: red shades – Lake Constance inflow populations, orange – Lake Constance littoral, and purple – Lake Constance pelagic population. Populations from the Baltic Sea (North-east Germany) are coded in blue, while North Sea samples are light blue (Norway and Great Britain); with the outgroup *G. nipponicus* shown in black. Bootstrap values calculated in percentage from all (n = 500) bootstrap replicates show confidence for each node, with exceptions where tree topology from the consensus tree was changed by TreeMix. Four was determined to be the optimum number of migration events according to model likelihoods. Arrows show the direction of admixture and are coloured according to migration weight, i.e., the proportion of ancestry received from the source population [0-0.5]. The level of genetic drift was indicated by branch length, with the scale bar showing 10 × the mean standard error of the sample covariance matrix. Matrices of the drift estimates and residuals are reported in Fig. S2.

### Genetic differentiation across the genome

To identify putative signatures of selection, we compared genetic differentiation between pelagic and littoral individuals across the genome. We detected 333 loci with *zF*_ST_*’* ≥ 3, distributed across the entire genome (Figure 4). Outlier loci showed on average increased absolute divergence (*D*_*xy*_) compared to the genomic background, both on autosomes (Wilcoxon: W = 2993041, p < 2.2e-16) and the sex chromosome (Wilcoxon: W = 6203, p = 0.01179) (Fig. 4B), suggesting that they are under divergent selection between littoral and pelagic sticklebacks. However, only 16 outlier loci showed strongly increased *D*_*xy*_ values (z*D*_*xy*_ ≥ 3). Comparisons between littoral and pelagic populations with inflow populations show a similar picture (Fig. S3 - S4).

**Fig. 4.**
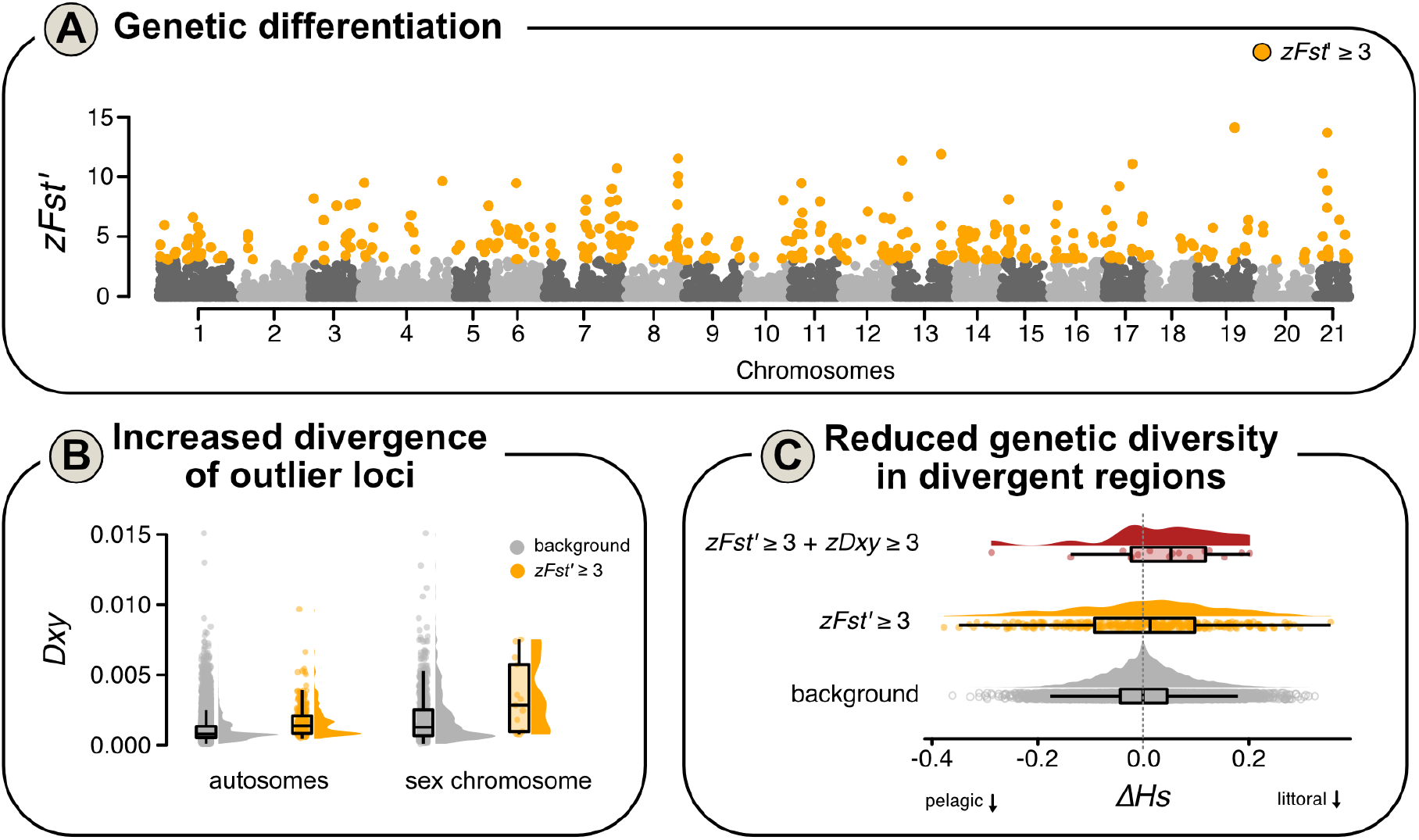
Signatures of selection. (**A**) Z-transformed F^ST^’ (zFst’) estimates for loci (dots) across all chromosomes (noted on the x-axis). Outlier loci with zF^ST^’ ≥ 3 are shown in orange. (**B**) Absolute divergence (Dxy) between outlier loci (orange) and the genomic background on autosomes and the sex chromosome. Individual dots denote genomic loci and the distribution of values is shown by density plots. The sex chromosome was analyses separately due to lower recombination rates compared to autosomes, and therefore potentially higher absolute divergence. (**C**) Comparison of delta gene diversity (ΔHs) between the genomic background (zFst’ < 3; grey), outlier loci (orange) and outlier loci showing increased absolute divergence (zDxy ≥ 3; red). Delta gene diversity was estimated by subtracting gene diversity in pelagic individuals from gene diversity in littoral individuals. Positive ΔHs values are indicative of reduced gene diversity in littoral individuals and vice versa. Individual dots denote genomic loci. Box plots defined in Fig. 2B.

We further tested for signals of divergent selection by comparing gene diversity (Fig. S5) between littoral and pelagic populations. While the mean between-population difference in gene diversity of outlier loci (*ΔHs*_*outlier*_) was not lower than the genomic background (*ΔHs*_*bg*_), outlier loci showed more extreme values than the genomic background (Kolmogorov-Smirnov test: D = 0.1694, p-value = 1.59e-08) (Fig. 4C). Outlier loci with extreme *ΔHs* values are more likely to be under divergent selection in the population with the lower gene diversity. We found that outlier loci with increased absolute divergence had on average lower gene diversity in littoral sticklebacks (positive *ΔH*_*S*_) (Fig. 4C), suggesting that these highly divergent loci are likely under selection in the littoral population.

### Marker associations with phenotypic traits

To identify genomic regions associated with ecologically-relevant phenotypes in Lake Constance stickleback, we performed genome-wide association analyses for later plate number and body shape using a BSLMM. For lateral plate number, 41 SNPs were associated (mean PIP > 0.01), with 7 SNPs (17.1%) showing very strong associations (mean PIP > 0.1). These were largely located on chromosome 4, and with a strongly associated SNP on chromosome 2 (Fig. 4A). For body shape, the BSLMM detected 104 associated SNPs with PIP > 0.01, and 1 strongly associated SNP on chromosome 21 with PIP > 0.1. Although body shape values were corrected for sex, a large proportion of associatied SNPs (n = 34; 32.7%) were located on the sex chromosome 19.

In line with the difference in the number of associated loci, estimates of the genetic architectures underlying lateral plate number and body shape, based on hyperparameters estimated using the BSLMM, revealed a similar picture (Fig. 5B). While the proportion of variance explained by all loci was similar for lateral plate number (PVE_*PN*_ = 88.4%) and body shape (PVE_*BS*_ = 85.3%), the lower proportion of PVE explained by sparse effect loci for body shape (PGE_*BS*_ = 61.5%, PGE_*PN*_ = 77.8%) and smaller estimated number of sparse effect loci for body shape compared to lateral plate number (mean *n gamma*_*BS*_ = 28; mean *n gamma*_*PN*_ = 6) supports a more polygenic basis of body shape.

**Fig. 5.**
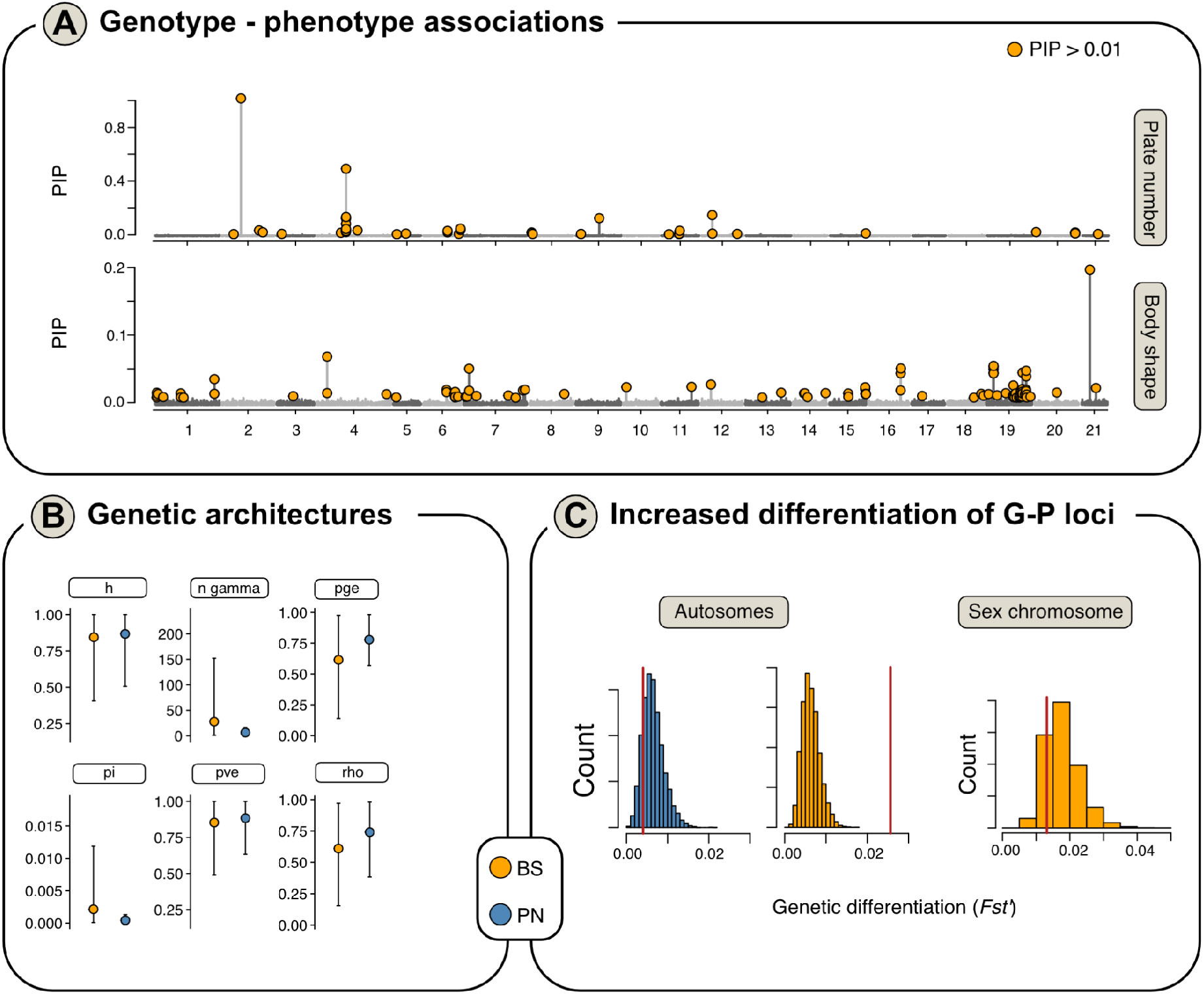
Genome-wide association analyses (GWAS) for phenotypic traits. (**A**) Posterior Inclusion Probabilities from BSLMMs for all SNPs (dots) across the genome are shown, with outliers SNPs passing significance threshold (PIP > 0.01) shown in red. Manhattan plots are shown for GWAS results with mean lateral plate number, body shape PC1-6, and total length. (**B**) Hyperparameters from BLSMMs are plotted, as the mean (large dot) and 95% confidence intervals (grey lines), for body shape (BS: yellow) and lateral plate number (PN: blue). (**C**) The distribution of genetic differentiation (F^ST^’) values for the permuted null distribution is shown as a histogram and mean differentiation for phenotype-associated loci on autosomes is indicated as a red line. Results are shown for SNPs associated with later plate number (blue) and body shape (yellow), for autosomes and the sex chromosome separately. No trait-associated SNPs were detected for later plate number on the sex chromosome.

To test if divergence in body shape between littoral and pelagic stickleback is driven by divergence of the underlying loci, we tested for genetic differentiation of these loci. Under selection on a polygenic genetic architecture (Fig. 3A), we might expect subtly increased genetic differentiation of many loci compared to the genomic background, rather than strong selection on a few loci. Indeed, body shape-associated loci were not strongly differentiated (i.e. outlier loci), but autosomal loci associated with body shape showed increased genetic differentiation between littoral and pelagic ecotypes compared to the genomic background (Fig. 5C). Loci associated with lateral plate number, which does not differ between pelagic and littoral individuals (Fig. 2B), did not show increased genetic differentiation compared to the genomic background. Body shape associated loci on the sex chromosomes also did not show increased genetic differentiation between littoral and pelagic individuals (Fig. 5C).

### Association of outlier loci with QTL for putatively adaptive traits

Of 1222 previously identified QTL involved in divergent phenotypic traits in three-spined sticklebacks (Peichel and Marques, 2017), 879 QTL (Supplementary Data S2) overlapped with *zF*_*ST*_’-outlier loci between pelagic and littoral populations. When considering only the strongest outlier loci, those with *zF*_*ST*_’ and *zDxy’* values equal or above three, we found 193 overlapping QTL regions. Feeding, body shape and defence related QTL were the most common outlier-associated QTL (n = 85, 67 and 19 QTL, respectively) (Fig. 6). The majority of the QTL explained little to intermediate amounts of variation (n = 173), while only one was a major effect locus explaining more than 20% of variation in total length.

**Fig. 6.**
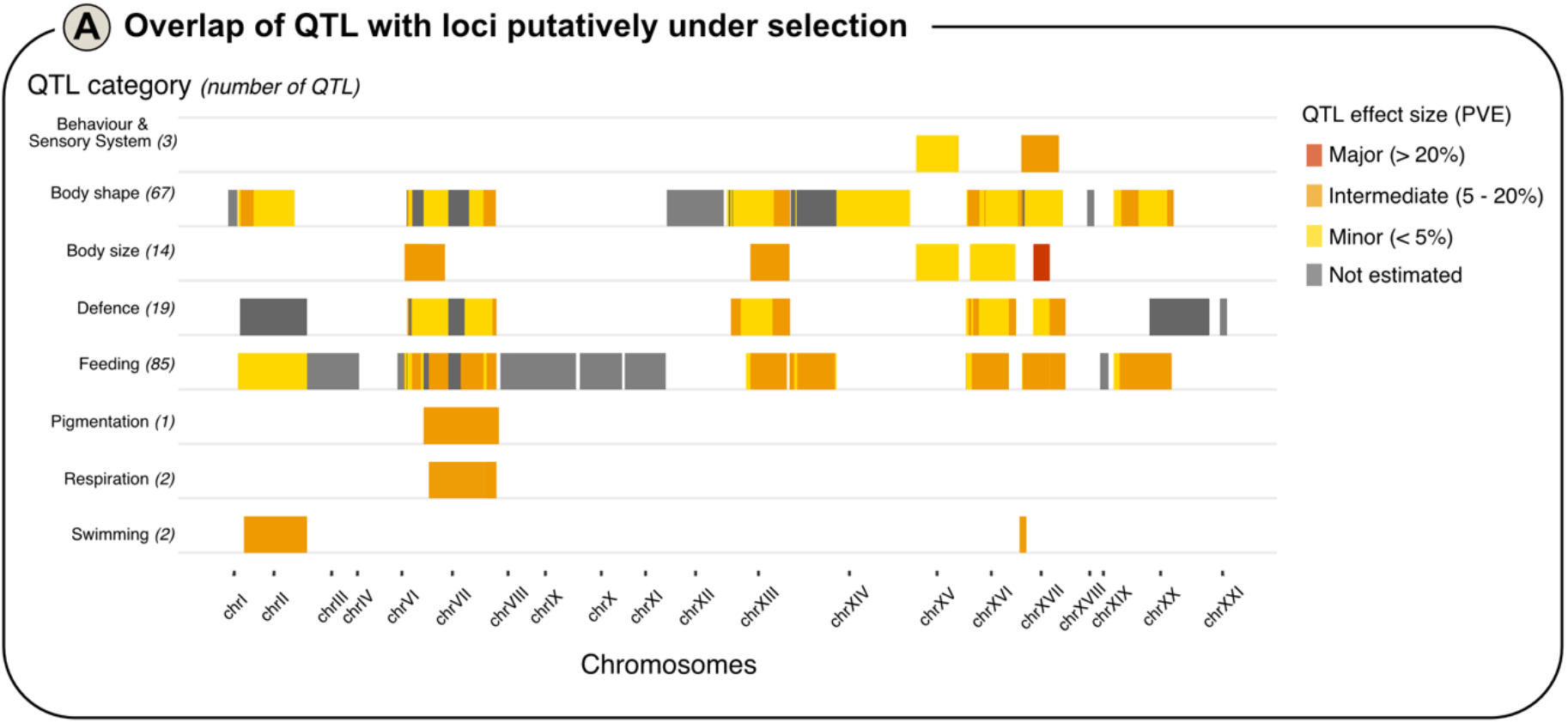
Genome-wide distribution of Quantitative Trait Loci among Lake Constance sticklebacks. Boxes show QTL (position along x-axis indicating chromosome number) that overlap with outlier loci (zFst’ ≥ 3 and zDxy ≥ 3) between pelagic and littoral stickleback. Effect sizes (minor, intermediate, major or not estimated) of trait variance are shown by their respective colour (see legend). Detailed positions are given in Supplementary Data S2.

Additionally, we identified numerous QTL (n = 351 for PIP ≥ 0.01; n = 159 for PIP ≥ 0.1) overlapping with loci associated with lateral plate number, including known QTL on chr.II and the major *EDA*-linked QTL on chr.IV (Peichel & Marques 2017) (Fig. S6A). The plate number QTL on chr.II, which were identified in ‘benthic x limnetic’ stickleback from Vancouver Island (Conte et al. 2015), overlapped with the locus that showed strongest association with lateral plate number in Lake Constance (PIP = 1). Similarly, many QTL (n = 551 for PIP ≥ 0.01; n = 29 for PIP ≥ 0.1) overlapped with body shape-associated loci (Fig. S6B). The single locus with the strongest association with body shape (PIP ≥ 0.1) co-localised with 29 QTL on chr.XXI that were largely associated with feeding, defence, and body shape-related traits, as well as swimming and neuromast patterning.

## Discussion

The aim of this study was to investigate the evolutionary origin, and phenotypic and genetic basis, of the recent appearance and rapid population increase of a pelagic ecotype of three-spined stickleback in Lake Constance. We found that the pelagic stickleback population in Lake Constance likely originated from the littoral population without any recent colonisation and/or introgression from other European stickleback lineages. Despite weak genome-wide divergence and high gene flow among lake populations, several genomic regions spread across the genome show increased genetic differentiation, partly co-localise with previously-identified QTL regions. Furthermore, we found that body shape-associated loci, a trait divergent between littoral and pelagic stickleback in Lake Constance, show increased genetic differentiation. This suggests that the rapid evolution and dramatic demographic expansion of the pelagic stickleback ecotype has been facilitated by large adaptive standing genetic diversity present within Lake Constance, rather than recently introduced adaptive genetic variation. Thus, effective management strategies have to focus on the entire stickleback population rather than the invasive pelagic ecotype.

### Phenotypic divergence of littoral and pelagic stickleback in Lake Constance

Sticklebacks are known to exhibit a high degree of phenotypic diversity between habitats, with some morphological traits evolving in parallel during their postglacial dispersal into new freshwater habitats (Bell and Foster 1994, McPhail 1993). In particular, the reduction of lateral plate armour in freshwater populations is regularly observed (Bell 2001, Hagen and Gilbertson 1971). Although this process can happen very quickly (Bell et al, 2004), our findings show that sticklebacks in Lake Constance still exhibit largely full armoring similar to their likely ancestral Baltic populations. Until the sudden mass abundance of the stickleback and their spread into the open water, the species was exclusively found in the areas close to the shore. The littoral fish community in Lake Constance includes several predatory fish species, including perch and pike, which are known to prey on sticklebacks (Jacobson *et al*., 2019, Donadi *et al*., 2017). Thus, we suggest that full plating may have remained relevant as effective predator protection (Reimchen 1994, Kitano *et al*., 2008, Rennison *et al*., 2019).

In contrast to lateral plate numbers, differences were found in body shape between habitats. In addition to general allometric effects and sexual dimorphism, there were significant differences between fish from the littoral zone compared to fish from the pelagic zone and river inflows (tributaries). Body shape analyses suggest that littoral and pelagic stickleback differ slightly in snout length and body depth, with longer snouts and deeper bodies in littoral fish. However, they do not strongly differ in trophic position (Gugele *et al*., submitted) and these differences are not very pronounced, and did not necessarily follow known patterns of divergence between benthic and limnetic sticklebacks in Canada (Schluter & McPhail 1992; Arnegard et al. 2014). However, future common garden experiments and more detailed phenotypic and trophic analyses could shed light on the eco-morphological basis of the rapid pelagic invasion of Lake Constance stickleback.

Habitat-specific morphological variation has already been shown in sticklebacks (Webster 2011, Gow *et al*., 2008) and differences in morphometric and meristic traits are found especially in systems with sympatric occurring limnetic and benthic populations (Nagel and Schluter 1998). In Lake Constance, studies on spatio-temporal movement of sticklebacks (Gugele *et al*., 2020), indicate temporal migration of sticklebacks from tributaries into the pelagic zone, and back. Recent studies on stable isotope ratios in the muscle and liver of sticklebacks could not find differences in trophic position between sticklebacks from pelagic and littoral habitats, but rather a mere preference to forage in the pelagic zone (Gugele *et al*., submitted), suggesting that body shape differences are potentially related to differences in habitat usage rather than trophic eco-morphology. Furthermore, size differences between sticklebacks from the pelagic and littoral zone may be explained by differences in sampling time, with fish from the pelagic zone being captured in late March, while fish from the littoral zone were captured in June and July. It can therefore be assumed that this is a time-dependent increase in size over the course of the year (Reimchen 1992, Yershov and Sukhotin 2015).

### Intralacustrine origin of pelagic stickleback in Lake Constance

To date, it has been unclear if pelagic stickleback in Lake Constance, which increased rapidly in abundance since 2012, originated within Lake Constance or are the result of a separate introduction (Eckmann & Engesser 2019). Our genetic results, based on PCA, admixture analyses, and phylogenetic comparison to other Northern European stickleback populations strongly suggest a recent intralacustrine origin of pelagic stickleback in Lake Constance. Genome-wide patterns of genetic differentiation were overall weak between habitats, suggesting either a very recent expansion into the pelagic by sticklebacks and/or ongoing gene flow. A very recent origin and ongoing gene flow are supported by annual hydroacoustic surveys (Eckmann & Engesser, 2019) and spatio-temporal sampling (Gugele *et al*., 2020), respectively.

The TreeMix analysis clustered stickleback from Lake Constance as sister to Baltic stickleback from Northern Germany, which is in line with the theory that Lake Constance was colonised by admixed populations which shared a substantial proportion of ancestry with marine-like sticklebacks repeatedly introduced to nearby streams from catchments south to the Baltic Sea around 150 years ago (Hudson *et al*., 2021; Muckle, 1972). Hybridization after secondary contact of deeply divergent North-eastern and Western European lineages has been argued to have facilitated rapid speciation in Lake Constance through provision of adaptive alleles (Marques et al. 2019; Hudson et al. 2021). However, we did not have the right samples in our study to test this hypothesis in respect to the littoral-pelagic divergence.

The slightly stronger clustering of pelagic and littoral stickleback in the PCA, and the lower genetic differentiation between littoral and pelagic sticklebacks, compared to the inflow populations might indicate that pelagic sticklebacks originated from the littoral population and that gene flow is higher between the littoral and pelagic zone compared to tributaries (Gugele *et al*., 2020). However, these genetic differences are very subtle and not sufficient to confirm putative spawning locations of pelagic stickleback. It has been hypothesised that pelagic sticklebacks migrate to tributaries to spawn. In general, the pelagic zone is an unusual habitat for three-spined sticklebacks, and thus not many such invasions, particularly in oligotrophic lakes, are known to date. Freshwater diversifications have mostly occurred along the lake-stream axis, and the very few cases of benthic-limnetic diversifications have been exclusive to small lakes in British Columbia, Canada (Baker *et al*., 2005; Schluter and McPhail, 1992) where sympatric ecotype evolution occurred over at least the past ten thousand years (Jones *et al*., 2012a). The latter emphasises the uniqueness of the contemporary pelagic invasion in Lake Constance. Unfortunately, the exact drivers for this behaviour still remain uncertain. Recently conducted studies could not find any evidence that parasite pressure or the use of planktonic food sources, which have not been exploited by pelagic fishes to date, are the cause of the expansion (Baer *et al*., 2022, Ogorelec *et al*., 2022). Furthermore, it has not been conclusively elucidated which role the potentially increased intraspecific competition in a growing littoral population since its establishment in the 1950’s (Hudson *et al*., 2021), as well as novel niche opportunities in the pelagic zone, such as opportunity for predation on naïve whitefish species (Ros *et al*., 2019; Roch *et al*., 2018), play in this process.

Overall, our study revealed the likely intralacustrine origin of pelagic Lake Constance sticklebacks from a single shared gene pool, yet the drivers of this rapid expansion and ecological diversification remain elusive.

### Polygenic basis of pelagic colonisation

Although genome-wide patterns of divergence suggest minimal genetic differentiation and ongoing gene flow, distinct genomic regions were differentiated between habitats and showed putative signs of selection. Despite the recent colonisation of the pelagic zone, we detected a polygenic signal of divergence with hundreds of outlier SNPs across the genome showing increased genetic differentiation between individuals from littoral and pelagic habitats. Genetic differentiation across large parts of the genome between benthic and limnetic sticklebacks was also observed in Canadian populations, despite an enrichment of divergent regions on certain chromosomes (Härer et al. 2021). In general, this genome-wide differentiation confirms the hypothesis that genetic differentiation is possible despite high connectivity over short evolutionary time periods. Such a scenario could have two non-mutually exclusive explanations: a) the ability of rapid adaptation of three-spined sticklebacks via sharing and reassembly of adaptive alleles through selection on standing genetic variation, which has been demonstrated to be a key mechanism in freshwater colonisation across numerous stickleback systems throughout its distribution range (e.g. Fang *et al*., 2020; Terekhanova *et al*., 2014); and b) strong divergent selection pressures arising from the differing environments in the pelagic *versus* littoral zone, which act on adaptive phenotypes such as body shape.

Three-spined stickleback genetic adaptation to novel environments in short time periods has been documented before, and similar explanations were proposed, such as in newly formed Alaskan ponds after an earthquake only 50 years ago (Lescak *et al*., 2015); as well as in Lake Constance, where selection signatures of lake-stream ecotype divergence was shown after only one generation (Laurentino *et al*., 2020). Similar rapid genetic differentiation from standing genetic variation over roughly 8 generations has also been documented in European whitefish following the re-oligotrophication of Lake Constance (Jacobs et al. 2020). In general, not only selective sweeps at a few loci but also polygenic responses to selection, as observed in our and other studies (Laurentino *et al*., 2020; Salmón et al. 2020), can lead to rapid local adaptation despite ongoing gene flow (Jain & Stephan 2017).

While genetic differentiation can occur without divergent selection, e.g. through linked selection in low recombination regions or genetic drift due to population bottlenecks, these are unlikely explanations in this system. Firstly, linked selection is less likely to lead to increased differentiation over such short evolutionary time scales (Burri 2017), and patterns of genetic diversity between habitats do not indicate the presence of genetic bottlenecks, which is supported by large observed stickleback populations (Eckmann & Engesser, 2019; Gugele et al. 2020). Furthermore, we observed increased absolute divergence in outlier loci, with strong divergence for a small subset of loci, suggesting a contribution of divergent selection in the genetic differentiation between habitats. Absolute divergence takes longer to build up than genetic differentiation, thus suggesting that increased divergence between pelagic and littoral sticklebacks is potentially due to the sorting of ancient adaptive alleles between habitats, which could have been potentially introduced through hybridization between stickleback lineages prior to the colonisation of Lake Constance (Marques et al. 2019; Hudson et al. 2021). However, differences in gene diversity at outlier loci between pelagic and littoral populations (Δ*Hs*_outlier_) compared to the genomic background, further indicate that genetic differentiation is potentially driven by divergent selection rather than variation in genetic diversity across the genome, which would be expected to lead to reduced diversity in both populations (Cruickshank & Hahn 2014; Burri 2017).

### Targets of selection

The identified genomic outlier loci raise the question whether these could be early signs of sympatric speciation in Lake Constance stickleback. In particular, the increased differentiation of body shape-associated autosomal loci between pelagic and littoral sticklebacks suggests that body shape, a trait that differs between populations in these habitats, is under divergent selection between habitats. This inference is furthermore strengthened by the fact that loci associated with lateral plate number, a trait that does not differ between pelagic and littoral sticklebacks, do not show increased differentiation. Thus, the observed signal is likely not due to chance. The genetic differentiation of body shape-associated loci furthermore suggests that the observed divergence between littoral and pelagic stickleback is not purely due to phenotypic plasticity but is at least partly genetically determined.

We did not test for genotype-association with variation in body size in our dataset, as stickleback were sampled at different timepoints throughout the year, even though divergence in body size in sticklebacks is a substantial driver of reproductive isolation as it affects reproductive behaviours such as mate choice (Moser *et al*., 2015) and thus may facilitate speciation in future generations. However, we found that differentiated loci, including those also showing strong absolute divergence, often co-localise with numerous and well-recognised quantitative trait loci for putatively adaptive traits, including body shape and body size, suggesting their role in phenotypic divergence. While respective outlier counts across trait categories could indicate relative relevance of traits under selection, biases could arise from disproportionate numbers of studies focusing on certain traits such as feeding (Peichel and Marques, 2017). Furthermore, the low density of RAD-seq derived SNPs across the genome makes it difficult to conclusively link SNPs to exact quantitative trait loci, as these SNPs are often likely not the target of selection and only linked to selected SNPs. Thus, to better understand the genomic basis of rapid phenotypic evolution in littoral-pelagic sticklebacks, more high-density genotyping or whole genome data will be needed.

## Conclusion

In conclusion, our results suggest that the novel pelagic *G. aculeatus* ecotype in Lake Constance, which has already caused ecosystem-wide effects on biodiversity and food-web integrity, likely arose within Lake Constance in less than 10 generations through sorting of standing genetic variation. Divergence in body shape between littoral and pelagic habitats, and potentially other relevant ecological and physiological phenotypic traits known from independent QTL studies, is reflective of divergent polygenic selection on trait-associated genes. However, the limited SNP-density across the genome precludes us from determining the most likely genomic targets of selection and phenotype-associated loci. Furthermore, more widespread outgroup genomes, particularly from the Western lineage that likely contributed adaptive alleles to the Lake Constance gene pool would be needed to more finely resolve the origin of alleles involved in the littoral-pelagic divergence. Common garden experiments and temporal phenotypic sampling in the wild could help to better understand the roles of evolutionary change versus plasticity in the rapid colonisation of the pelagic zone and identify putatively adaptive phenotypic traits. A better understanding of the processes facilitating the rapid invasion of the pelagic zone of Lake Constance, which had drastic ecosystem-wide effects, could facilitate management of this population and also in other systems with rapid pelagic invasions, such as the Baltic Sea. Our results suggest that the observed pelagic colonisation likely resulted through a combination of the adaptive capabilities of three-spined sticklebacks through large standing genetic variation and a so far unknown change in ecosystem properties, which needs to be identified for the implementation of effective control measures and to be able to predict future occurrences of such invasion events. In the end, our results highlight that the entire stickleback population, and no specific sub-population, is potentially capable of employing the pelagic zone, or can re-invade the pelagic zone, and thus needs to be managed as a whole.

## Acknowledgements

Special thanks to Sarah Gugele, Jan Baer and Andreas Revermann, who planned and carried out the sampling. Sarah Gugele also performed the initial phenotyping. Thanks to Cameron Hudson for the organisational support with shipping the genetic samples. Thanks to Jonas Reiter for the assistance in staining the fish and conducting the meristic counts. Sam Fenton and Darren Hunter are thanked for helping with coding aspects of the analyses. Thank you to Jasminca Behrmann-Godel without whom this collaboration would not have taken place.

This study was supported by the grant “SeeWandel: Life in Lake Constance - the past, present and future” within the framework of the Interreg V programme “Alpenrhein-Bodensee-Hochrhein (Germany/Austria/Switzerland/Liechtenstein)” which funds are provided by the European Regional Development Fund as well as the Swiss Confederation and cantons. Arne Jacobs was supported by a Leverhulme Trust Early Career Fellowship (ECF-2020-509) and a University of Glasgow Lord Kelvin/Adam Smith Leadership Fellowship. The funders had no role in study design, data collection and analysis, decision to publish, or preparation of the manuscript.

## Data Archiving Statement

The raw RAD-seq data in this study are available in the NCBI SRA under the BioProject XXXXXXX with the following run accessions: SRRXXX. Phenotypic data and the filtered VCF file with SNPs are deposited in the University of Glasgow Enlighten repository (DOIXXXXX). [Data will be made available upon acceptance of the manuscript].

## Author contributions

A.B. conceived the project. Sampling was performed by A.R., and phenotypic analysis was performed by S.R. Genomic analysis was performed by C.D and A.J, with input from K.R.E. C.D. and S.R. wrote the first version of the manuscript with input from A.J. All authors contributed to the final version.

## Supplementary Material

Supplementary Datasets S1 and S2 as appendix.

***Supplementary Data S1: Sample information***. *Information on species, sample location, population, sample IDs, ecotype, geographic coordinates, and citations for studies from which samples were re-acquired. Samples were either whole genome sequences (WGS) or reduced representation sequences (RAD-seq, recognition site enzyme SbfI)*.

***Supplementary Data S2: List of Quantitative Trait Loci***. *Listed are 287 QTLs where matches with outlier loci in Lake Constance were identified. Information on position of QTL (chromosome, start and end position in base pair number), as well as the associated phenotypic trait and trait category is given. Percent Variance Explained (PVE) of the respective traits by QTLs was grouped into three categories: minor (<5%), intermediate (∼5-20%), major ∼>20%) or was otherwise noted as undetermined. For each study which identified respective QTLs, relevant population and ecotype information of three-spined stickleback populations used in the analyses are provided. These QTLs originate from a list of 1034 QTLs (685 retained in this study) compiled by Peichel and Marques (2017)*.

**Supplementary Table S1.** Results of the permutation-based Procrustes ANOVA testing influence of different factors on stickleback shape. Df = degrees of freedom, SS = sums of squares, MS = mean square, Rsq = R-squared, F = F-Value, Z = effect size, P = p-value.

| Factor | Df | SS | MS | Rsq | F | Z | P-value |
| --- | --- | --- | --- | --- | --- | --- | --- |
| Log(size) | 1 | 0.017447 | 0.0174465 | 0.13950 | 21.6642 | 5.5338 | < 0.0001* |
| Sex | 1 | 0.019460 | 0.0194599 | 0.15560 | 24.1644 | 6.2986 | < 0.0001* |
| Sex:site | 8 | 0.022123 | 0.0027654 | 0.17689 | 3.4339 | 5.8960 | < 0.0001* |
| Residuals | 82 | 0.066036 | 0.0008053 | 0.52801 |  |  |  |

\*statistically significant ( $\alpha = 0.05$ )

**Supplementary Table S2.** Results of the permutation-based Procrustes ANOVA testing for common or unique allometric effects. ResDf = residual degrees of freedom, Df = degrees of freedom, SS = sums of squares, MS = mean square, Rsq = R-squared, F = F-Value, Z = effect size, P = p-value.

| Model | ResDf | Df | SS | MS | Rsq | F | Z | P-value |
| --- | --- | --- | --- | --- | --- | --- | --- | --- |
| Coords ~ log(size) + sex/site | 82 | 1 |  |  | 0 |  |  |  |
| Coords ~ log(size) * sex/site | 81 | 1 | 0.0009105 | 0.000911 | 0.00728 | 1.132 | 0.512 | 0.285 |

\*statistically significant ( $\alpha = 0.05$ )

**Supplementary Table S3:** Genome-wide genetic divergence. Genetic differentiation (F_st_’ / upper triangle) and absolute genetic divergence (D_xy_ / lower triangle) between all sampled populations.

| $F_{st}'/D_{xy}$ | BRU | Pelagic | ALT | Littoral | NON |
| --- | --- | --- | --- | --- | --- |
| BRU |  | 0.00571 | 0.00839 | 0.00483 | 0.00851 |
| Pelagic | 0.00117 |  | 0.00378 | 0.00189 | 0.00456 |
| ALT | 0.00118 | 0.00113 |  | 0.00426 | 0.00706 |
| Littoral | 0.00116 | 0.00111 | 0.00112 |  | 0.00426 |
| NON | 0.00117 | 0.00112 | 0.00114 | 0.00111 |  |

**Supplementary Table S4:** Support for migration edges. Four migration events inferred by TreeMix have a source (the set of populations placed in the subtree below the origin of the migration edge) and recipient (the group of populations placed in the subtree below the destination of the migration edge). For each migration event the mean weight of the migration (w)̅, i.e., the proportion of ancestry received by the donor population, the mean jackknife estimate of the weight 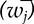 and the jackknife estimate of the standard error 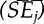 averaged over 30 independent runs (N), as well as the least significant p-value recovered from all runs are indicated.

| Source | Recipient | $\bar{w}$ | $\bar{w}_j$ | $\overline{SE}_j$ | p-value | N |
| --- | --- | --- | --- | --- | --- | --- |
| NOR_R | NOR_L | 0.3178 | 0.3176 | 0.0278 | <0.0001 | 30 |
| *G. nipponicus* | PEL | 0.0279 | 0.0279 | 0.0117 | 0.0085 | 30 |
| TY_M | GER_R, GER_L | 0.1258 | 0.1259 | 0.0465 | 0.0034 | 30 |
| PEL | *G. nipponicus* | 0.0912 | 0.0912 | 0.0191 | <0.0001 | 30 |

**Supplementary Figure S1:**
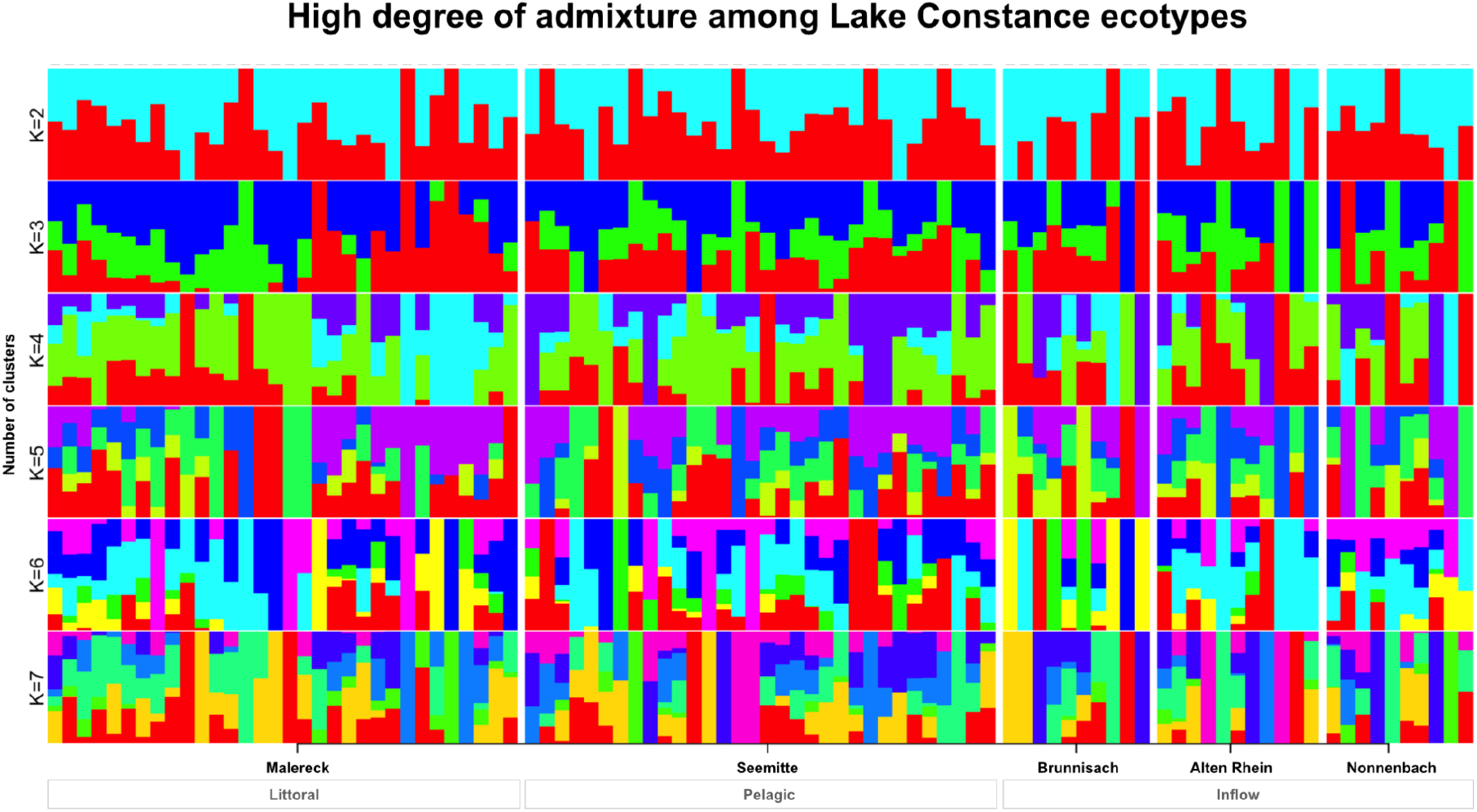
Admixture analysis across Lake Constance three-spined stickleback ecotypes. Genetic structure was best explained with one genetic cluster (K=1) based on ten-fold cross-validation error. Shown are results for analyses with 2-7 clusters showing random admixture for each individual across all populations which are grouped by ecotype (littoral, pelagic and inflow). Analysis was performed on pruned data excluding the sex chromosome (∼11 thousand SNPs).

**Supplementary Figure S2:**
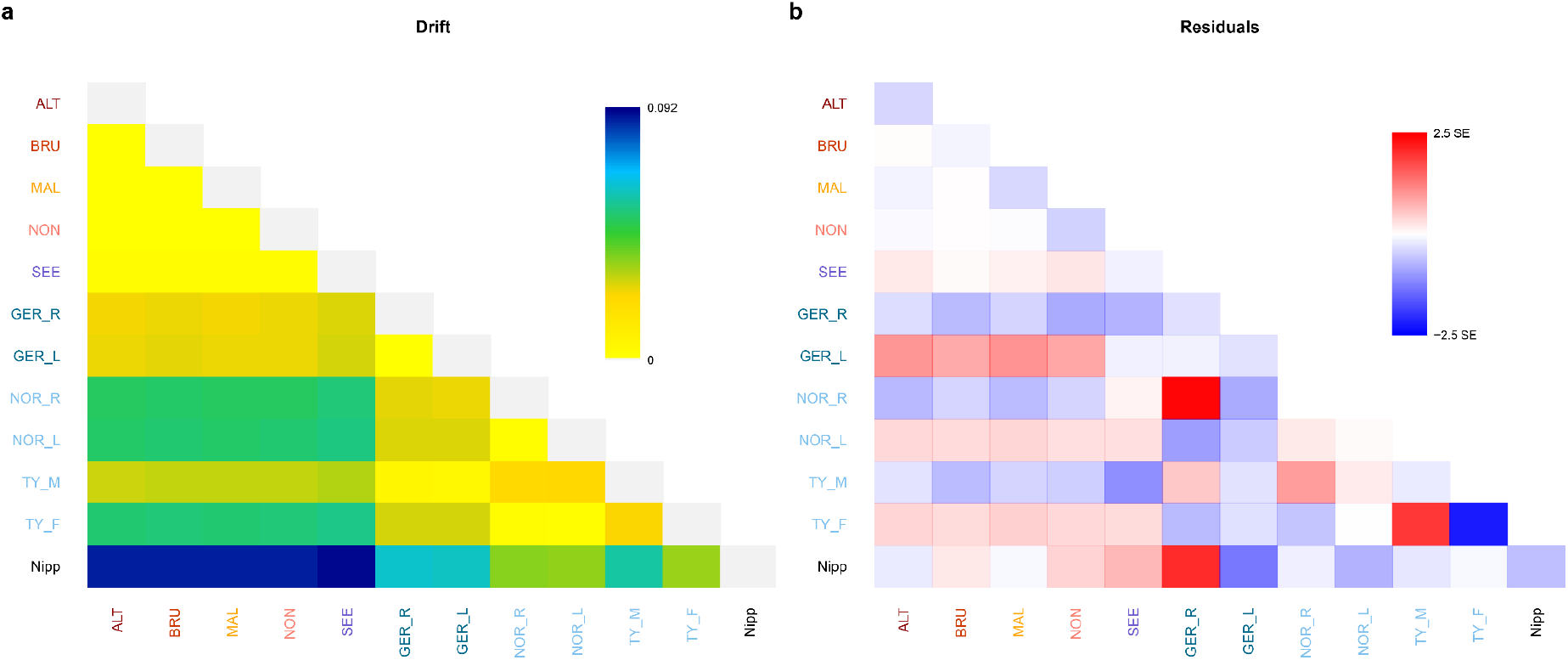
Residuals and drift estimates from Maximum likelihood tree. **a, Pairwise drift** estimates from the ML tree (Fig. 8) calculated under a Gaussian model by TreeMix. Populations are coloured according to location: red shades – Lake Constance inflow populations, orange – Lake Constance littoral, and purple – Lake Constance pelagic population. Population outgroups are coloured in blue (Baltic Sea), and light blue (North Sea). **b, Pairwise residuals** from the ML tree (Fig. 8) with colour coding as above. Positive residuals suggest closer relationships between populations than as inferred by the tree, while negative residuals indicate more distant relationships.

**Supplementary Figure S3:**
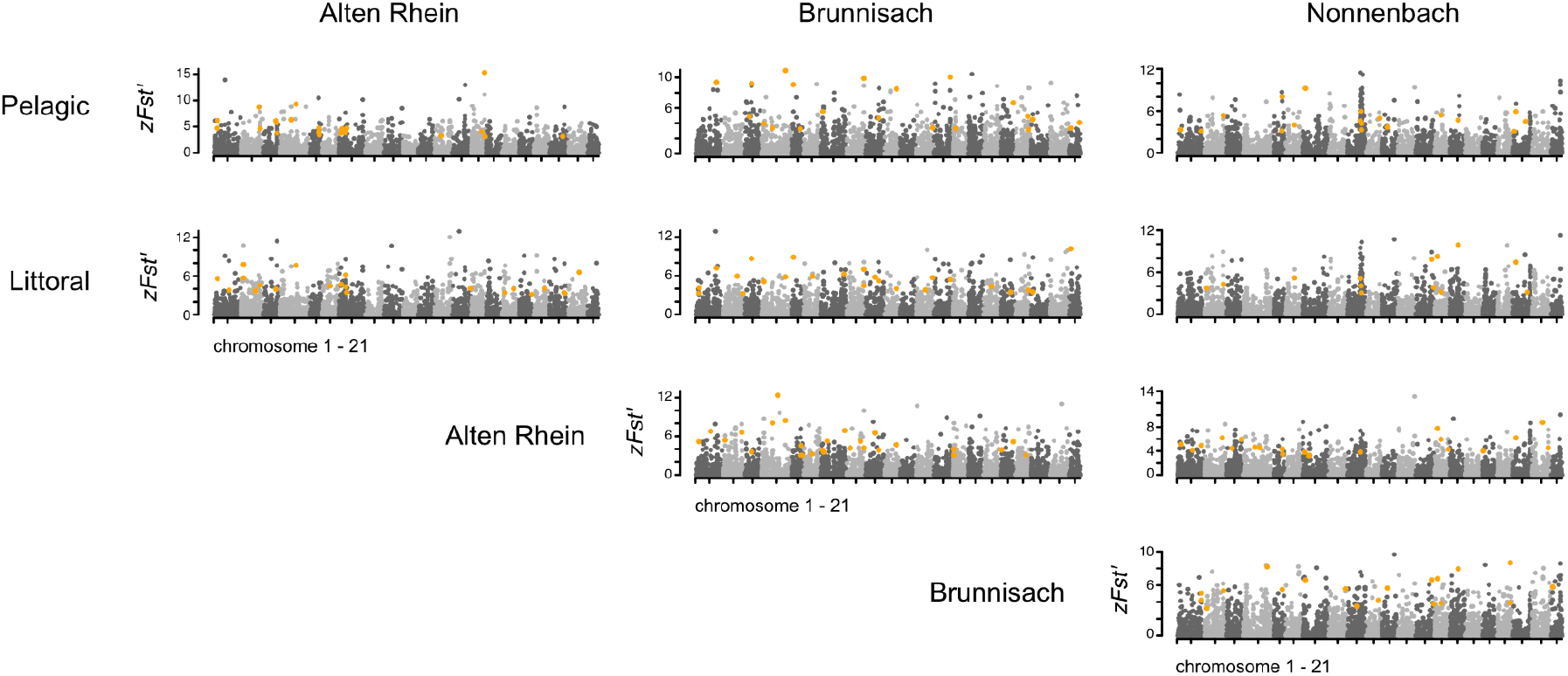
Manhattan plots of genome-wide genetic differentiation. Z-transformed F_ST_’ estimates for loci (dots) across all chromosomes (noted on the x-axis) are shown. Loci with z-transformed D_xy_ ≥ 3 are highlighted in orange.

**Supplementary Figure S4:**
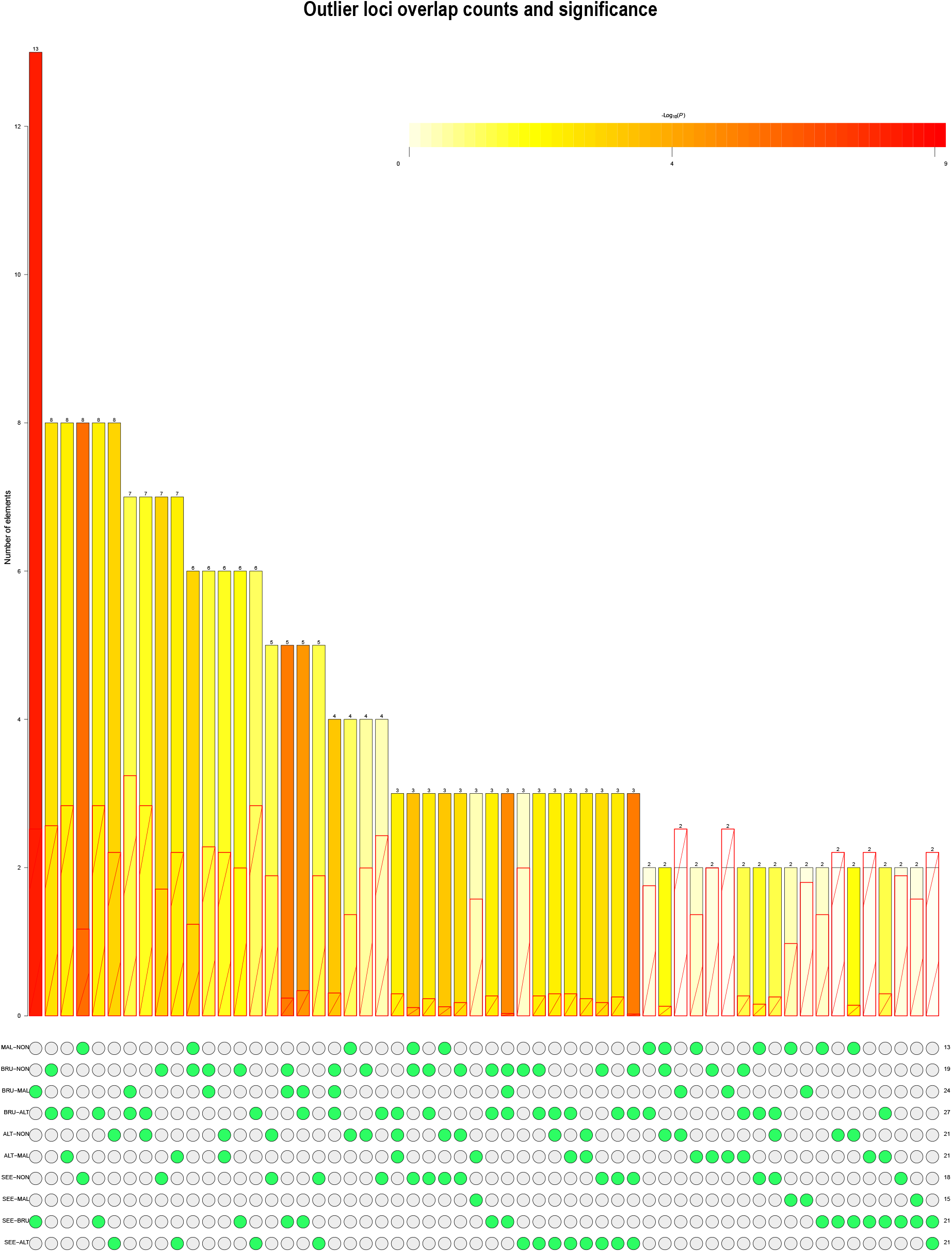
Overlaps of outlier loci from all pair-wise population comparisons. The observed number of outlier loci overlaps (along the y-axis) for different sets (green points along the x-axis) of all combinations of pairwise comparisons including inflow (ALT, BRU, NON), littoral (MAL) and pelagic (SEE) populations. Outlier loci for each population pair were those with z-transformed F_ST_’ ≥ 3 and zDxy ≥ 3. The colour scale of log-transformed p-values indicates whether the observed count of overlaps was significantly higher than the expected number (red bars). Shown are only those sets with at least 2 overlaps. Calculations and plotting was carried out in the SuperExactTest R Package v.1.0.7 (Wang et al., 2015).

**Supplementary Figure S5:**
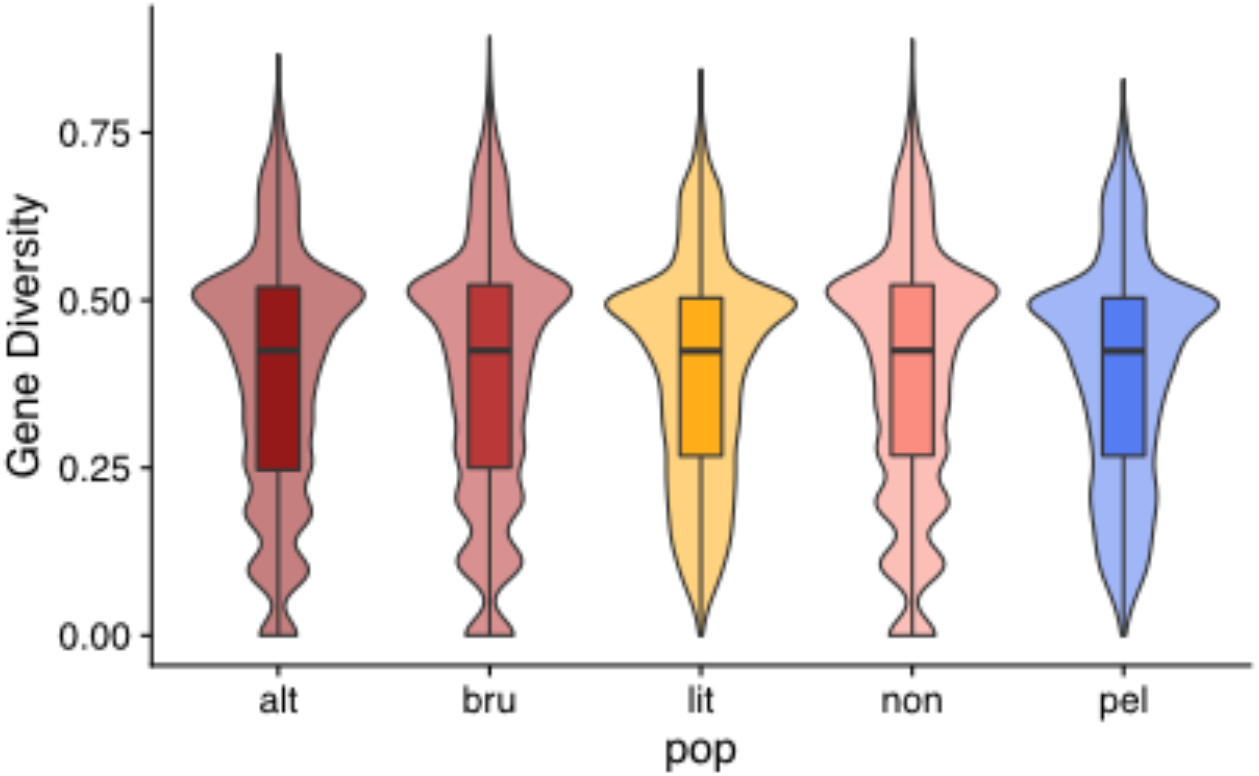
Patterns of gene diversity across populations. Haplotype-based gene diversity across all loci is plotted for each sampling site/population as violin plots and overlaid boxplots.

**Supplementary Figure S6:**
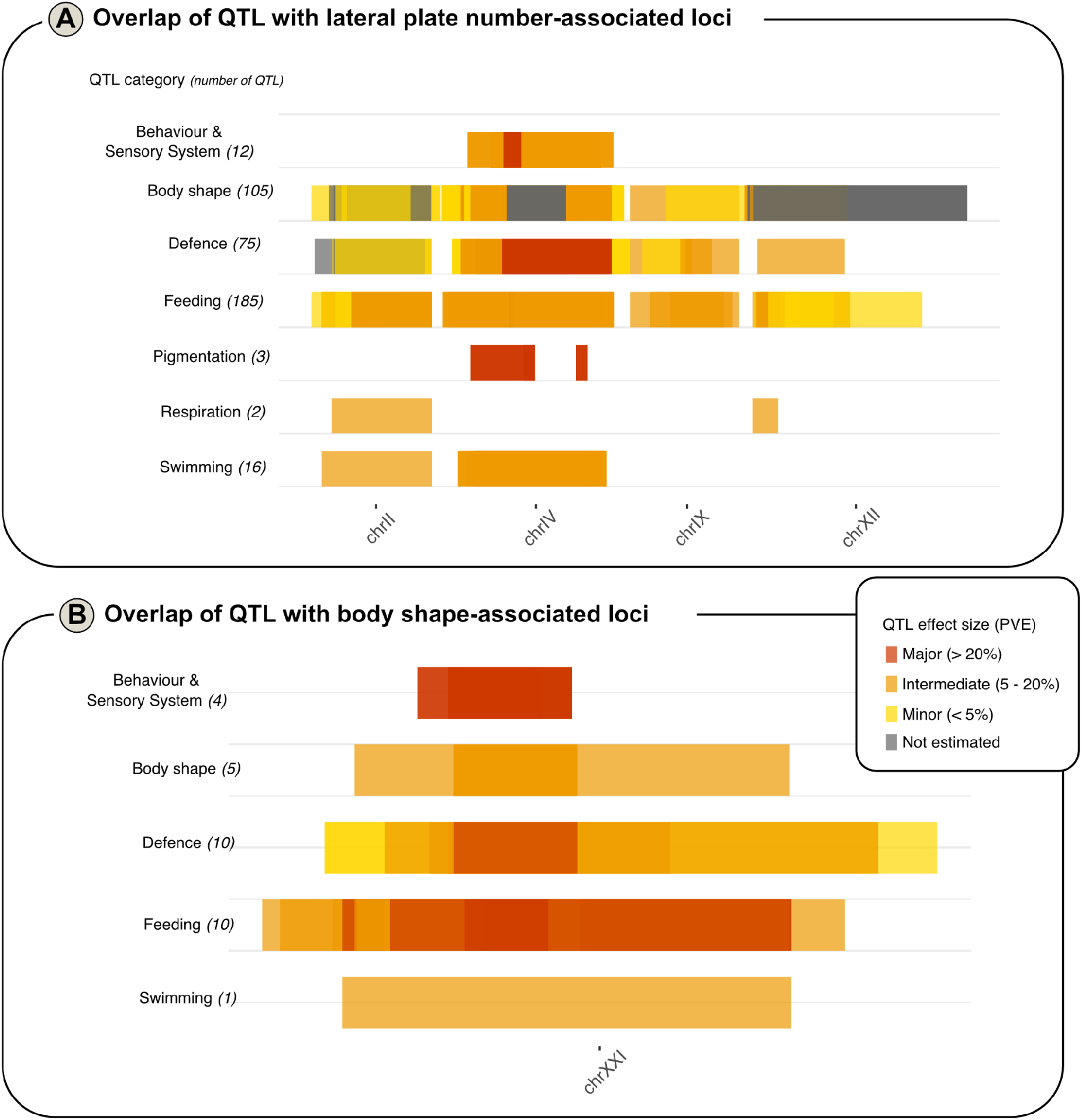
QTL overlap for loci associated with plate number and body shape. (**A-B**) Boxes indicate QTL positions (along x-axis indicating chromosome number) and effect sizes (minor, intermediate, major or not estimated) of trait variance explained by their respective colour. Shown are QTL overlapping with loci strongly associated (PIP > 0.1) with (A) lateral plate number and (B) body shape. Only chromosomes with overlapping QTL are shown.

## Notes

### Competing Interest Statement

The authors have declared no competing interest.

## References

Albert, A.Y., Sawaya, S., Vines, T.H., Knecht, A.K., Miller, C.T., Summers, B.R., Balabhadra, S., Kingsley, D.M., & Schluter, D. (2008). The genetics of adaptive shape shift in stickleback: pleiotropy and effect size. Evolution: International Journal of Organic Evolution, 62(1), 76–85.

Albert, J.S., Destouni, G., Duke-Sylvester, S.M., Magurran, A.E., Oberdorff, T., Reis, R.E., Winemiller, K.O., & Ripple, W.J. (2021). Scientists’ warning to humanity on the freshwater biodiversity crisis. Ambio, 50(1), 85–94.

Alexander, D.H., Novembre, J., & Lange, K. (2009). Fast model-based estimation of ancestry in unrelated individuals. Genome Research, 19(9), 1655–1664.

Alexander, T., Vonlanthen, P., Périat, G., Raymond, J., Degiorgi, F., & Seehausen, O. (2016). Artenvielfalt und Zusammensetzung der Fischpopulation im Bodensee. Kastanienbaum: Projet Lac, Eawag.

Armstrong, C., Richardson, D. S., Hipperson, H., Horsburgh, G. J., Küpper, C., Percival-Alwyn, L., … & Spurgin, L. G. (2018). Genomic associations with bill length and disease reveal drift and selection across island bird populations. Evolution Letters, 2(1), 22–36.

Arnegard, M.E., McGee, M.D., Matthews, B., Marchinko, K.B., Conte, G.L., Kabir, S., Bedford, N., Bergek, S., Chan, Y.F., & Jones, F.C. (2014). Genetics of ecological divergence during speciation. Nature, 511(7509), 307–311.

Baer, J., Eckmann, R., Rösch, R., Arlinghaus, R., & Brinker, A. (2017). Managing Upper Lake Constance fishery in a multi-sector policy landscape: beneficiary and victim of a century of anthropogenic trophic change. Inter-sectoral governance of inland fisheries. Too Big To Ignore-WorldFish, St. John’s, Newfoundland, Canada, 32–47.

Baer, J., Gugele, S. M., Bretzel, J., DeWeber, J. T., & Brinker, A. (2021). All day-long: Sticklebacks effectively forage on whitefish eggs during all light conditions. Plos One, 16(8), e0255497.

Baken, E. K., Collyer, M. L., Kaliontzopoulou, A., & Adams, D. C. (2021). geomorph v4. 0 and gmShiny: Enhanced analytics and a new graphical interface for a comprehensive morphometric experience. Methods in Ecology and Evolution, 12(12), 2355–2363.

Baker, J.A., Cresko, W.A., Foster, S.A., & Heins, D.C. (2005). Life-history differentiation of benthic and limnetic ecotypes in a polytypic population of threespine stickleback (Gasterosteus aculeatus). Evolutionary Ecology Research, 7(1), 121–131.

Bax, N., Williamson, A., Aguero, M., Gonzalez, E., & Geeves, W. (2003). Marine invasive alien species: a threat to global biodiversity. Marine Policy, 27(4), 313–323.

Bell, M. A., & Foster, S. A. (1994). The evolutionary biology of the threespine stickleback. Oxford University Press, Oxford, UK.

Bell, M. A. (2001). Lateral plate evolution in the threespine stickleback: getting nowhere fast. In Microevolution Rate, Pattern, Process, 445–461. Springer, Dordrecht.

Bell, M. A., Aguirre, W. E., & Buck, N. J. (2004). Twelve years of contemporary armor evolution in a threespine stickleback population. Evolution, 58(4), 814–824.

Bergström, U., Olsson, J., Casini, M., Eriksson, B.K., Fredriksson, R., Wennhage, H., & Appelberg, M. (2015). Stickleback increase in the Baltic Sea–a thorny issue for coastal predatory fish. Estuarine, Coastal and Shelf Science, 163, 134–142.

Berner, D. (2021). Re-evaluating the evidence for facilitation of stickleback speciation by admixture in the Lake Constance basin. Nature Communications, 12, 2806.

Berner, D., Moser, D., Roesti, M., Buescher, H., & Salzburger, W. (2014). Genetic architecture of skeletal evolution in European lake and stream stickleback. Evolution 68(6), 1792–1805.

Bretzel, J.B., Geist, J., Gugele, S.M., Baer, J., & Brinker, A. (2021). Feeding Ecology of Invasive Three-Spined Stickleback (Gasterosteus aculeatus) in Relation to Native Juvenile Eurasian Perch (Perca fluviatilis) in the Pelagic Zone of Upper Lake Constance. Frontiers in Environmental Science, 9, 254.

Burri, R. (2017) Linked selection, demography and the evolution of correlated genomic landscapes in birds and beyond. Molecular Ecology, 26, 3853–3856.

Byström, P., Bergström, U., Hjälten, A., Ståhl, S., Jonsson, D., & Olsson, J. (2015). Declining coastal piscivore populations in the Baltic Sea: Where and when do sticklebacks matter? Ambio, 44(3), 462–471.

Donadi, S., Austin, Å. N., Bergström, U., Eriksson, B. K., Hansen, J. P., Jacobson, P., & Eklöf, J. S. (2017). A cross-scale trophic cascade from large predatory fish to algae in coastal ecosystems. Proceedings of the Royal Society B: Biological Sciences, 284(1859), 20170045.

Catchen, J., Hohenlohe, P. A., Bassham, S., Amores, A., & Cresko, W. A. (2013). Stacks: an analysis tool set for population genomics. Molecular Ecology, 22(11), 3124–3140.

Charlesworth, B. (1998). Measures of divergence between populations and the effect of forces that reduce variability. Molecular Biology and Evolution, 15(5), 538–543.

Charlesworth, B., Nordborg, M., & Charlesworth, D. (1997). The effects of local selection, balanced polymorphism and background selection on equilibrium patterns of genetic diversity in subdivided populations. Genetics Research, 70(2), 155–174.

Chaves, J. A., Cooper, E. A., Hendry, A. P., Podos, J., De León, L. F., Raeymaekers, J. A., … & Uy, J. A. C. (2016). Genomic variation at the tips of the adaptive radiation of Darwin’s finches. Molecular Ecology, 25(21), 5282–5295.

Cleves, P.A., Ellis, N.A., Jimenez, M.T., Nunez, S.M., Schluter, D., Kingsley, D.M., & Miller, C.T. (2014). Evolved tooth gain in sticklebacks is associated with a cis-regulatory allele of Bmp6. Proceedings of the National Academy of Sciences 111(38), 13912–13917.

Collyer, M. L., & Adams, D. C. (2018). RRPP: An r package for fitting linear models to high-dimensional data using residual randomization. Methods in Ecology and Evolution, 9(7), 1772–1779.

Collyer, M. L. & Adams, D. C. (2021). RRPP: Linear Model Evaluation with Randomized Residuals in a Permutation Procedure, R package version 1.1.2. https://cran.r-project.org/package=RRPP.

Colosimo, P.F., Peichel, C.L., Nereng, K., Blackman, B.K., Shapiro, M.D., Schluter, D., Kingsley, D.M., & Patel, N. (2004). The genetic architecture of parallel armor plate reduction in threespine sticklebacks. PLoS Biology, 2(5), e109.

Comeault, A. A., Soria-Carrasco, V., Gompert, Z., Farkas, T. E., Buerkle, C. A., Parchman, T. L., & Nosil, P. (2014). Genome-wide association mapping of phenotypic traits subject to a range of intensities of natural selection in Timema cristinae. The American Naturalist, 183(5), 711–727.

Conte, G.L., Arnegard, M.E., Best, J., Chan, Y.F., Jones, F.C., Kingsley, D.M., Schluter, D., & Peichel, C.L. (2015). Extent of QTL reuse during repeated phenotypic divergence of sympatric threespine stickleback. Genetics, 201(3), 1189–1200.

Copp, G., Bianco, P., Bogutskaya, N.G., Erős, T., Falka, I., Ferreira, M.T., Fox, M.G., Freyhof, J., Gozlan, R., & Grabowska, J. (2005). To be, or not to be, a non-native freshwater fish? Journal of Applied Ichthyology, 21(4), 242–262.

Cox, J.G., & Lima, S.L. (2006). Naiveté and an aquatic–terrestrial dichotomy in the effects of introduced predators. Trends in Ecology & Evolution, 21(12), 674–680.

Coyle, S.M., Huntingford, F.A., & Peichel, C.L. (2007). Parallel evolution of Pitx1 underlies pelvic reduction in Scottish threespine stickleback (Gasterosteus aculeatus). Journal of Heredity, 98(6), 581–586.

Cresko, W.A., Amores, A., Wilson, C., Murphy, J., Currey, M., Phillips, P., Bell, M.A., Kimmel, C.B., & Postlethwait, J.H. (2004). Parallel genetic basis for repeated evolution of armor loss in Alaskan threespine stickleback populations. Proceedings of the National Academy of Sciences, 101(16), 6050–6055.

Crooks, J.A. (2002). Characterizing ecosystem-level consequences of biological invasions: the role of ecosystem engineers. Oikos, 97(2), 153–166.

Cruickshank, T.E., & Hahn, M.W. (2014). Reanalysis suggests that genomic islands of speciation are due to reduced diversity, not reduced gene flow. Molecular Ecology, 23(13), 3133–3157.

Cucherousset, J., & Olden, J.D. (2011). Ecological impacts of nonnative freshwater fishes. Fisheries, 36(5), 215–230.

Dahms, C., Kemppainen, P., Zanella, L.N., Zanella, D., Carosi, A., Merilä, J., & Momigliano, P. (2022). Cast away in the Adriatic: Low degree of parallel genetic differentiation in three-spined sticklebacks. Molecular Ecology, 31(4), 1234–1253.

Danecek, P., Auton, A., Abecasis, G., Albers, C.A., Banks, E., DePristo, M.A., Handsaker, R.E., Lunter, G., Marth, G.T., & Sherry, S.T. (2011). The variant call format and VCFtools. Bioinformatics, 27(15), 2156–2158.

Darwall, W., Bremerich, V., De Wever, A., Dell, A.I., Freyhof, J., Gessner, M.O., Grossart, H.P., Harrison, I., Irvine, K., & Jähnig, S.C. (2018). The Alliance for Freshwater Life: a global call to unite efforts for freshwater biodiversity science and conservation. Aquatic Conservation: Marine and Freshwater Ecosystems, 28(4), 1015–1022.

DeWeber, J. T., Baer, J., Rösch, R. & Brinker, A. (2022). Turning summer into winter: nutrient dynamics, temperature, density dependence and invasive species drive bioenergetic processes and growth of a keystone coldwater fish. Oikos, e09316.

Dudgeon, D. (2019). Multiple threats imperil freshwater biodiversity in the Anthropocene. Current Biology, 29(19), R960–R967.

Eckmann, R., & Engesser, B. (2019). Reconstructing the build-up of a pelagic stickleback (Gasterosteus aculeatus) population using hydroacoustics. Fisheries Research, 210, 189–192.

Eckmann, R., Gerster, S., & Kraemer, A. (2006). Yields of European perch from Upper Lake Constance from 1910 to present. Fisheries Management and Ecology, 13(6), 381–390.

Eklöf, J.S., Sundblad, G., Erlandsson, M., Donadi, S., Hansen, J.P., Eriksson, B.K., & Bergström, U. (2020). A spatial regime shift from predator to prey dominance in a large coastal ecosystem. Communications Biology, 3(1), 1–9.

Ellis, N.A., Glazer, A.M., Donde, N.N., Cleves, P.A., Agoglia, R.M., & Miller, C.T. (2015). Distinct developmental genetic mechanisms underlie convergently evolved tooth gain in sticklebacks. Development, 142(14), 2442–2451.

Erickson, P.A., Glazer, A.M., Killingbeck, E.E., Agoglia, R.M., Baek, J., Carsanaro, S.M., Lee, A.M., Cleves, P.A., Schluter, D., & Miller, C.T. (2016). Partially repeatable genetic basis of benthic adaptation in threespine sticklebacks. Evolution, 70(4), 887–902.

Etter, P.D., Preston, J.L., Bassham, S., Cresko, W.A., & Johnson, E.A. (2011). Local de novo assembly of RAD paired-end contigs using short sequencing reads. PloS One, 6(4), e18561.

Fang, B., Merilä, J., Matschiner, M., & Momigliano, P. (2020). Estimating uncertainty in divergence times among three-spined stickleback clades using the multispecies coalescent. Molecular Phylogenetics and Evolution, 142, 106646.

Fang, B., Merilä, J., Ribeiro, F., Alexandre, C.M., & Momigliano, P. (2018). Worldwide phylogeny of three-spined sticklebacks. Molecular Phylogenetics and Evolution, 127, 613–625.

Felsenstein, J. (2005). PHYLIP (Phylogeny Inference Package). Version 3.6.

Feulner, P.G., Chain, F.J., Panchal, M., Huang, Y., Eizaguirre, C., Kalbe, M., Lenz, T.L., Samonte, I.E., Stoll, M., & Bornberg-Bauer, E. (2015). Genomics of divergence along a continuum of parapatric population differentiation. PLoS Genetics, 11(2), e1004966.

Fitak, R.R. (2018). (Submitted). optM: an R package to optimize the number of migration edges using threshold models. Journal of Heredity.

Gallardo, B., Clavero, M., Sánchez, M.I., & Vilà, M. (2016). Global ecological impacts of invasive species in aquatic ecosystems. Global Change Biology, 22(1), 151–163.

Glazer, A.M., Cleves, P.A., Erickson, P.A., Lam, A.Y., & Miller, C.T. (2014). Parallel developmental genetic features underlie stickleback gill raker evolution. Evodevo, 5(1), 1–16.

Glazer, A. M., Killingbeck, E. E., Mitros, T., Rokhsar, D. S., & Miller, C. T. (2015). Genome assembly improvement and mapping convergently evolved skeletal traits in sticklebacks with genotyping-by-sequencing. G3: Genes, Genomes, Genetics, 5(7), 1463–1472.

Goodall, C. (1991). Procrustes methods in the statistical analysis of shape. Journal of the Royal Statistical Society: Series B (Methodological), 53(2), 285–321.

Gow, J. L., Rogers, S. M., Jackson, M., & Schluter, D. (2008). Ecological predictions lead to the discovery of a benthic–limnetic sympatric species pair of threespine stickleback in Little Quarry Lake, British Columbia. Canadian Journal of Zoology, 86(6), 564–571.

Gower, J. C. (1975). Generalized procrustes analysis. Psychometrika, 40(1), 33–51.

Greenwood, A. K., Ardekani, R., McCann, S. R., Dubin, M. E., Sullivan, A., Bensussen, S., … & Peichel, C. L. (2015). Genetic mapping of natural variation in schooling tendency in the threespine stickleback. G3: Genes, Genomes, Genetics, 5(5), 761–769.

Greenwood, A. K., Jones, F. C., Chan, Y. F., Brady, S. D., Absher, D. M., Grimwood, J., … & Peichel, C. L. (2011). The genetic basis of divergent pigment patterns in juvenile threespine sticklebacks. Heredity, 107(2), 155–166.

Greenwood, A. K., Wark, A. R., Yoshida, K., & Peichel, C. L. (2013). Genetic and neural modularity underlie the evolution of schooling behavior in threespine sticklebacks. Current Biology, 23(19), 1884–1888.

Gude, H., & Gries, T. (1998). Phosporus fluxes in Lake Constance (with 12 figures and 4 tables). Ergebnisse der Limnologie, (53), 505–544.

Gugele, S. M., Baer, J., & Brinker, A. (2020). The spatiotemporal dynamics of invasive three-spined sticklebacks in a large, deep lake and possible options for stock reduction. Fisheries Research, 232, 105746.

Gugele, S. M., Baer, J., Spießl, C., Yohannes, E., Blumenshine, S., & Brinker, A. (submitted). Stable isotope values and trophic position of a unique invasion of the three-spined stickleback in Lake Constance indicates significant piscivory.

Hagen, D. W., & Gilbertson, L. G. (1972). Geographic variation and environmental selection in Gasterosteus aculeatus L. in the Pacific Northwest, America. Evolution, 32–51.

Härer, A., Bolnick, D. I., & Rennison, D. J. (2021). The genomic signature of ecological divergence along the benthic-limnetic axis in allopatric and sympatric threespine stickleback. Molecular Ecology, 30(2), 451–463.

Herten, K., Hestand, M. S., Vermeesch, J. R., & Van Houdt, J. K. (2015). GBSX: a toolkit for experimental design and demultiplexing genotyping by sequencing experiments. BMC Bioinformatics, 16(1), 1–6.

Holm, S. (1979). A simple sequentially rejective multiple test procedure. Scandinavian Journal of Statistics, 65–70.

Hudson, C. M., Lucek, K., Marques, D. A., Alexander, T. J., Moosmann, M., Spaak, P., … & Matthews, B. (2021). Threespine stickleback in lake constance: The ecology and genomic substrate of a recent invasion. Frontiers in Ecology and Evolution, 8, 611672.

Hunt, S.E., McLaren, W., Gil, L., Thormann, A., Schuilenburg, H., Sheppard, D., Parton, A., Armean, I.M., Trevanion, S.J., & Flicek, P. (2018). Ensembl variation resources. Database 2018.

Jacobs, Arne, et al. (2019) Rapid niche expansion by selection on functional genomic variation after ecosystem recovery. Nature Ecology & Evolution, 3(1), 77–86.

Jakobsson, M., Edge, M. D., & Rosenberg, N. A. (2013). The relationship between F ST and the frequency of the most frequent allele. Genetics, 193(2), 515–528.

Jacobson, P., Bergström, U., & Eklöf, J. (2019). Size-dependent diet composition and feeding of Eurasian perch (Perca fluviatilis) and northern pike (Esox lucius) in the Baltic Sea.

Jain, K., & Stephan, W. (2017). Modes of rapid polygenic adaptation. Molecular Biology and Evolution, 34(12), 3169–3175.

Jones, F.C., Chan, Y.F., Schmutz, J., Grimwood, J., Brady, S.D., Southwick, A.M., Absher, D.M., Myers, R.M., Reimchen, T.E., & Deagle, B.E. (2012a). A genome-wide SNP genotyping array reveals patterns of global and repeated species-pair divergence in sticklebacks. Current Biology, 22(1), 83–90.

Jones, F.C., Grabherr, M.G., Chan, Y.F., Russell, P., Mauceli, E., Johnson, J., Swofford, R., Pirun, M., Zody, M.C., & White, S. (2012b). The genomic basis of adaptive evolution in threespine sticklebacks. Nature, 484(7392), 55–61.

Kimmel, C. B., Ullmann, B., Walker, C., Wilson, C., Currey, M., Phillips, P. C., … & Cresko, W. A. (2005). Evolution and development of facial bone morphology in threespine sticklebacks. Proceedings of the National Academy of Sciences, 102(16), 5791–5796.

Kiruba-Sankar, R., Raj, J. P., Saravanan, K., Kumar, K. L., Angel, J. R. J., Velmurugan, A., & Roy, S. D. (2018). Invasive species in freshwater ecosystems–threats to ecosystem services. In Biodiversity and climate change adaptation in Tropical islands (pp. 257–296). Academic Press.

Kitano, J., Ross, J.A., Mori, S., Kume, M., Jones, F.C., Chan, Y.F., Absher, D.M., Grimwood, J., Schmutz, J., & Myers, R.M. (2009). A role for a neo-sex chromosome in stickleback speciation. Nature, 461(7267), 1079–1083.

Kleinlercher, G., Muerth, P., Pohl, H., & Ahnelt, H. (2008). Welche Stichlingsart kommt in Österreich vor, Gasterosteus aculeatus oder Gasterosteus gymnurus. Österreichs Fischerei, 61, 158–161.

Knapp, R. A., & Matthews, K. R. (2000). Non-native fish introductions and the decline of the mountain yellow-legged frog from within protected areas. Conservation Biology, 14(2), 428–438.

Korneliussen, T. S., Albrechtsen, A. & Nielsen, R. (2014). ANGSD: analysis of next generation sequencing data. BMC Bioinformatics, 15(1), 1–13.

Laurentino, T. G., Moser, D., Roesti, M., Ammann, M., Frey, A., Ronco, F., … & Berner, D. (2020). Genomic release-recapture experiment in the wild reveals within-generation polygenic selection in stickleback fish. Nature communications, 11(1), 1–9.

Leinonen, T., Herczeg, G., Cano, J. M., & Merilä, J. (2011). Predation-imposed selection on threespine stickleback (Gasterosteus aculeatus) morphology: a test of the refuge use hypothesis. Evolution: International Journal of Organic Evolution, 65(10), 2916–2926.

Le Rouzic, A., Østbye, K., Klepaker, T. O., Hansen, T. F., Bernatchez, L., Schluter, D., & Vøllestad, L. A. (2011). Strong and consistent natural selection associated with armour reduction in sticklebacks. Molecular Ecology, 20(12), 2483–2493.

Lescak, E. A., Bassham, S. L., Catchen, J., Gelmond, O., Sherbick, M. L., von Hippel, F. A., & Cresko, W. A. (2015). Evolution of stickleback in 50 years on earthquake-uplifted islands. Proceedings of the National Academy of Sciences, 112(52), E7204–E7212.

Li, H., Durbin, R. (2010). Fast and accurate long-read alignment with Burrows–Wheeler transform. Bioinformatics, 26(5), 589–595.

Li, H., Handsaker, B., Wysoker, A., Fennell, T., Ruan, J., Homer, N., Marth, G., Abecasis, G., & Durbin, R. (2009). The sequence alignment/map format and SAMtools. Bioinformatics, 25(16), 2078–2079.

Liu, J., Shikano, T., Leinonen, T., Cano, J. M., Li, M. H., & Merilä, J. (2014). Identification of major and minor QTL for ecologically important morphological traits in three-spined sticklebacks (Gasterosteus aculeatus). G3: Genes, Genomes, Genetics, 4(4), 595–604.

Lowe, W. H., Muhlfeld, C. C., & Allendorf, F. W. (2015). Spatial sorting promotes the spread of maladaptive hybridization. Trends in Ecology & Evolution, 30(8), 456–462.

Lucek, K., Lemoine, M., & Seehausen, O. (2014a). Contemporary ecotypic divergence during a recent range expansion was facilitated by adaptive introgression. Journal of Evolutionary Biology, 27(10), 2233–2248.

Lucek, K., Sivasundar, A., Kristjánsson, B. K., Skúlason, S., & Seehausen, O. (2014b). Quick divergence but slow convergence during ecotype formation in lake and stream stickleback pairs of variable age. Journal of Evolutionary Biology, 27(9), 1878–1892.

Lymbery, A. J., Morine, M., Kanani, H. G., Beatty, S. J., & Morgan, D. L. (2014). Co-invaders: the effects of alien parasites on native hosts. International Journal for Parasitology: Parasites and Wildlife, 3(2), 171–177.

McPhail, J. D. (1993). Ecology and evolution of sympatric sticklebacks (Gasterosteus): origin of the species pairs. Canadian Journal of Zoology, 71(3), 515–523.

Mainka, S. A., & Howard, G. W. (2010). Climate change and invasive species: double jeopardy. Integrative Zoology, 5(2), 102–111.

Malek, T. B., Boughman, J. W., Dworkin, I. A. N., & Peichel, C. L. (2012). Admixture mapping of male nuptial colour and body shape in a recently formed hybrid population of threespine stickleback. Molecular Ecology, 21(21), 5265–5279.

Marques, D. A., Lucek, K., Meier, J. I., Mwaiko, S., Wagner, C. E., Excoffier, L., & Seehausen, O. (2016). Genomics of rapid incipient speciation in sympatric threespine stickleback. PLoS Genetics, 12(2), e1005887.

Marques, D. A., Lucek, K., Sousa, V. C., Excoffier, L., & Seehausen, O. (2019). Admixture between old lineages facilitated contemporary ecological speciation in Lake Constance stickleback. Nature communications, 10(1), 1–14.

Miller, C. T., Beleza, S., Pollen, A. A., Schluter, D., Kittles, R. A., Shriver, M. D., & Kingsley, D. M. (2007). cis-Regulatory changes in Kit ligand expression and parallel evolution of pigmentation in sticklebacks and humans. Cell, 131(6), 1179–1189.

Miller, C. T., Glazer, A. M., Summers, B. R., Blackman, B. K., Norman, A. R., Shapiro, M. D., … & Kingsley, D. M. (2014). Modular skeletal evolution in sticklebacks is controlled by additive and clustered quantitative trait loci. Genetics, 197(1), 405–420.

Moser, D., Kueng, B., & Berner, D. (2015). Lake-stream divergence in stickleback life history: a plastic response to trophic niche differentiation?. Evolutionary Biology, 42(3), 328–338.

Moyle, P. B., & Light, T. (1996). Biological invasions of fresh water: empirical rules and assembly theory. Biological conservation, 78(1-2), 149–161.

Muckle, R. (1972). Der dreistachlige Stichling (Gasterosteus aculeatus L.) im Bodensee. Schriften des Vereins für Geschichte des Bodensees und seiner Umgebung.

Muhlfeld, C. C., Kalinowski, S. T., McMahon, T. E., Taper, M. L., Painter, S., Leary, R. F., & Allendorf, F. W. (2009). Hybridization rapidly reduces fitness of a native trout in the wild. Biology letters, 5(3), 328–331.

Nagel, L., & Schluter, D. (1998). Body size, natural selection, and speciation in sticklebacks. Evolution, 52(1), 209–218.

Nei, M. (1973). Analysis of gene diversity in subdivided populations. Proceedings of the National Academy of Sciences, 70(12), 3321–3323.

Nei, M. (1987). Molecular evolutionary genetics. Columbia University Press.

Ogorelec, Ž., Rudstam, L. G., & Straile, D. (2022). Can young-of-the-year invasive fish keep up with young-of-the-year native fish? A comparison of feeding rates between invasive sticklebacks and whitefish. Ecology and Evolution, 12(1), e8486.

Peichel, C. L., & Marques, D. A. (2017). The genetic and molecular architecture of phenotypic diversity in sticklebacks. Philosophical Transactions of the Royal Society B: Biological Sciences, 372(1713), 20150486.

Peichel, C. L., Nereng, K. S., Ohgi, K. A., Cole, B. L., Colosimo, P. F., Buerkle, C. A., … & Kingsley, D. M. (2001). The genetic architecture of divergence between threespine stickleback species. Nature, 414(6866), 901–905.

Peichel, C. L., Ross, J. A., Matson, C. K., Dickson, M., Grimwood, J., Schmutz, J., … & Kingsley, D. M. (2004). The master sex-determination locus in threespine sticklebacks is on a nascent Y chromosome. Current Biology, 14(16), 1416–1424.

Peters, R. H. (1986). The ecological implications of body size. Cambridge University Press, Cambridge.

Petri, M. (2006). Water quality of lake constance. The Rhine, 127–138.

Phillips, B. L., & Perkins, T. A. (2019). Spatial sorting as the spatial analogue of natural selection. Theoretical Ecology, 12(2), 155–163.

Pickrell, J., & Pritchard, J. (2012). Inference of population splits and mixtures from genome-wide allele frequency data. Nature Precedings, 1–1.

Purcell, S., Neale, B., Todd-Brown, K., Thomas, L., Ferreira, M. A., Bender, D., et. al. (2007). PLINK: a tool set for whole-genome association and population-based linkage analyses. The American Journal of Human Genetics, 81(3), 559–575.

Reimchen, T. E. (1992). Extended longevity in a large-bodied stickleback, Gasterosteus, population. Canadian field-naturalist. Ottawa ON, 106(1), 122–125.

Ricciardi, A., & Rasmussen, J. B. (1999). Extinction rates of North American freshwater fauna. Conservation biology, 13(5), 1220–1222.

Roch, S., von Ammon, L., Geist, J., & Brinker, A. (2018). Foraging habits of invasive three-spined sticklebacks (Gasterosteus aculeatus)–impacts on fisheries yield in Upper Lake Constance. Fisheries Research, 204, 172–180.

Roesti, M., Moser, D., & Berner, D. (2013). Recombination in the threespine stickleback genome—patterns and consequences. Molecular ecology, 22(11), 3014–3027.

Rogers, S. M., Tamkee, P., Summers, B., Balabahadra, S., Marks, M., Kingsley, D. M., & Schluter, D. (2012). Genetic signature of adaptive peak shift in threespine stickleback. Evolution: International Journal of Organic Evolution, 66(8), 2439–2450.

Rösch, R., Baer, J., & Brinker, A. (2018). Impact of the invasive three-spined stickleback (Gasterosteus aculeatus) on relative abundance and growth of native pelagic whitefish (Coregonus wartmanni) in Upper Lake Constance. Hydrobiologia, 824(1), 243–254.

Ros, A., Dunst, J., Gugele, S., & Brinker, A. (2019). Anti-predator mechanisms in evolutionarily predator-naïve vs. adapted fish larvae. Ecosphere, 10(4), e02699.

Salmón, P., Jacobs, A., Ahrén, D., Biard, C., Dingemanse, N. J., Dominoni, D. M., … & Isaksson, C. (2021). Continent-wide genomic signatures of adaptation to urbanisation in a songbird across Europe. Nature Communications, 12(1), 1–14.

Schluter, D., & McPhail, J. D. (1992). Ecological character displacement and speciation in sticklebacks. The American Naturalist, 140(1), 85–108.

Schmid, D. W., McGee, M. D., Best, R. J., Seehausen, O., & Matthews, B. (2019). Rapid divergence of predator functional traits affects prey composition in aquatic communities. The American Naturalist, 193(3), 331–345.

Shapiro, M. D., Marks, M. E., Peichel, C. L., Blackman, B. K., Nereng, K. S., Jónsson, B., … & Kingsley, D. M. (2004). Genetic and developmental basis of evolutionary pelvic reduction in threespine sticklebacks. Nature, 428(6984), 717–723.

Sharpe, D. M., Räsänen, K., Berner, D., & Hendry, A. P. (2008). Genetic and environmental contributions to the morphology of lake and stream stickleback: implications for gene flow and reproductive isolation. Evolutionary Ecology Research, 10(6), 849–866.

Shine, R., Brown, G. P., & Phillips, B. L. (2011). An evolutionary process that assembles phenotypes through space rather than through time. Proceedings of the National Academy of Sciences, 108(14), 5708–5711.

Sokal, R.R., and Rohlf, J.F. (2003). Biometry: The Principles and Practice of Statistics in Biological Research. W.H. Freemann and Company, New York, USA.

Spence, R., Wootton, R. J., Barber, I., Przybylski, M., & Smith, C. (2013). Ecological causes of morphological evolution in the three-spined stickleback. Ecology and Evolution, 3(6), 1717–1726.

Steel, R. G. (1960). A rank sum test for comparing all pairs of treatments. Technometrics 2, 197–207.

Stich, H. B., & Brinker, A. (2010). Oligotrophication outweighs effects of global warming in a large, deep, stratified lake ecosystem. Global Change Biology, 16(2), 877–888.

Strayer, D. L. (2010). Alien species in fresh waters: ecological effects, interactions with other stressors, and prospects for the future. Freshwater Biology, 55, 152–174.

Team, R. (2019). RStudio: Integrated Development for R (RStudio, Inc.).

Team, R.C. (2013). R: A language and environment for statistical computing.

Terekhanova, N. V., Logacheva, M. D., Penin, A. A., Neretina, T. V., Barmintseva, A. E., Bazykin, G. A., … & Mugue, N. S. (2014). Fast evolution from precast bricks: genomics of young freshwater populations of threespine stickleback Gasterosteus aculeatus. PLoS Genetics, 10(10), e1004696.

Wang, M., Zhao, Y., & Zhang, B. (2015). Efficient test and visualization of multi-set intersections. Scientific Reports, 5(1), 1–12.

Wark, A. R., Mills, M. G., Dang, L. H., Chan, Y. F., Jones, F. C., Brady, S. D., … & Peichel, C. L. (2012). Genetic architecture of variation in the lateral line sensory system of threespine sticklebacks. G3: Genes, Genomes, Genetics, 2(9), 1047–1056.

Werner, S., Bauer, H.-G., Heine, G., Jacoby, H., and Stark, H. (2018). 55 Jahre Wasservogelzählung am Bodensee: Bestandsentwicklung der Wasservögel von 1961/62 bis 2015/16.

Yershov, P., & Sukhotin, A. (2015). Age and growth of marine three-spined stickleback in the White Sea 50 years after a population collapse. Polar Biology, 38(11), 1813–1823.

Yong, L., Peichel, C. L., & McKinnon, J. S. (2016). Genetic architecture of conspicuous red ornaments in female threespine stickleback. G3: Genes, Genomes, Genetics, 6(3), 579–588.

Yoshida, K., Makino, T., Yamaguchi, K., Shigenobu, S., Hasebe, M., Kawata, M., … & Kitano, J. (2014). Sex chromosome turnover contributes to genomic divergence between incipient stickleback species. PLoS Genetics, 10(3), e1004223.

Zhou, X., Carbonetto, P., & Stephens, M. (2013). Polygenic modeling with Bayesian sparse linear mixed models. PLoS Genetics, 9(2), e1003264.

